# Flexible categorization in perceptual decision making

**DOI:** 10.1101/2020.05.23.110460

**Authors:** Genís Prat-Ortega, Klaus Wimmer, Alex Roxin, Jaime de la Rocha

## Abstract

Perceptual decisions require the brain to make categorical choices based on accumulated sensory evidence. The underlying computations have been studied using either phenomenological drift diffusion models or neurobiological network models exhibiting winner-take-all attractor dynamics. Although both classes of models can account for a large body of experimental data, it remains unclear to what extent their dynamics are qualitatively equivalent. Here we show that, unlike the drift diffusion model, the attractor model can operate in different integration regimes: an increase in the stimulus fluctuations or the stimulus duration promotes transitions between decision-states leading to a crossover between weighting mostly early evidence (primacy regime) to weighting late evidence (recency regime). Between these two limiting cases, we found a novel regime, which we name *flexible categorization*, in which fluctuations are strong enough to reverse initial categorizations, but only if they are incorrect. This asymmetry in the reversing probability results in a non-monotonic psychometric curve, a novel and distinctive feature of the attractor model. Finally, we show psychophysical evidence for the crossover between integration regimes predicted by the attractor model and for the relevance of this new regime. Our findings point to correcting transitions as an important yet overlooked feature of perceptual decision making.

## Introduction

Integrating information over time is a fundamental computation that neural systems can adaptively perform in a variety of contexts. The integration of perceptual evidence is an example of such computation, and its most common paradigm is the binary categorization of ambiguous stimuli characterized by a stream of sensory evidence. This process is typically modeled with the drift diffusion model with absorbing bounds (DDMA) which integrates the stimulus evidence linearly until one of the bounds is reached ^1^. The DDMA and its different variations have been successfully used to fit psychometric and chronometric curves ^2^, to capture the speed accuracy trade off ^1,3,4^, to account for history dependent choice biases ^5^, changes of mind ^6^, confidence reports ^7^ or the Weber’s law ^8^. Although the absorbing bounds were originally thought of as a mechanism to terminate the integration process, the DDMA has also been applied to fixed duration tasks ^9,10^. In motion discrimination tasks, for instance, it can reproduce the subjects’ tendency to give more weight to early rather than late stimulus information, which is called a primacy effect ^9,11–15^. However, depending on the details of the task and the stimulus, subjects can also give more weight to late rather than to early evidence (i.e. a recency effect) ^16,17^ or weigh the whole stimulus uniformly ^18^. In order to account for these differences, the DDMA needs to be modified by using reflecting instead of absorbing bounds or by removing the bounds altogether ^19^. Despite their considerable success in fitting experimental data, the DDMA and its many variants remain purely phenomenological descriptions of sensory integration. This makes it difficult to link the DDMA to the actual neural circuit mechanisms underlying perceptual decision making.

These neural circuit mechanisms have been studied with biophysical attractor network models that can integrate stimulus evidence over relatively long time scales ^20,21^. Attractor network models have been used, among other examples, to study the adjustment of speed-accuracy trade-off in a cortico-basal ganglia circuit ^22^, learning dynamics of sensorimotor associations ^23^, the generation of choice correlated sensory activity in hierarchical networks ^24,25^, the role of the pulvino-cortical pathway in controlling the effective connectivity within and across cortical regions ^26^ or how trial history biases can emerge from the circuit dynamics ^27^. In the typical regime in which the attractor network was originally used for perceptual categorization ^20^, the impact of the stimulus on the decision decreases as the network evolves towards an attractor. In this regime, the integration dynamics of the attractor model are qualitatively similar to those of the DDMA as it also shows a primacy effect. Moreover, the attractor network can also provide an excellent fit to psychometric and chronometric curves ^20,28^. Thus, a common implicit assumption is that the attractor network is simply a neurobiological implementation of the DDMA ^29,30^ and hence there has been more interest in studying the similarities between these two models rather than their differences ^31^ (but see ^32,33^).

Here, we show that the attractor model has richer dynamics beyond the well known primacy regime. In particular, the model exhibits a crossover from primacy to recency as the stimulus fluctuations or stimulus duration are increased. Intermediate to these two limiting regimes, the stimulus can impact the upcoming decision nearly uniformly across the entire stimulus duration. Specifically, if the first attractor state reached corresponds to the incorrect choice, stimulus fluctuations later in the trial can lead to a correcting transition, while if the initial attractor is correct, fluctuations are not strong enough to drive an error transition. As a consequence, the model shows a non-monotonic psychometric curve as a function of the strength of stimulus fluctuations, and the maximum occurs precisely in this intermediate “flexible categorization” regime. To illustrate the relevance of our theoretical results, we re-analyze data from two psychophysical experiments ^34,35^ and show that the attractor model can quantitatively fit the crossover from primacy to recency with the stimulus duration, and the integration and storage of evidence when stimuli are separated by a memory delay. Our characterization of the flexible categorization regime in the attractor model reveals that correcting transitions may be a key property of evidence integration in perceptual decision-making.

## Results

### Canonical models of perceptual decision making show invariant dynamics

We start by characterizing the dynamics of evidence integration in standard drift-diffusion models during a binary classification task. These models are described as the evolution of a decision variable *x(t)* that integrates the moment-by-moment evidence *S(t)* provided by the stimulus, plus a noise term reflecting the internal stochasticity in the process ^1,29,31^: 

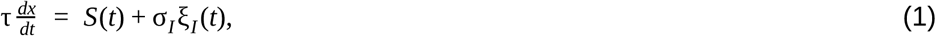

where τ is the time constant of the integration process. The evidence *S*(*t*) fluctuates in time and can be written as a constant mean drift μ, plus a time-varying term, caused by the fluctuations of the input stimulus: *S*(*t*) = μ + σ_*S*_ ξ_*S*_ (*t*). We call *σ*_*S*_ the magnitude of stimulus fluctuations. Assuming that both fluctuating terms, ξ_*I*_ and ξ_*S*_ are Gaussian stochastic processes, Equation 1 can be recast as the motion of a particle in a potential: 

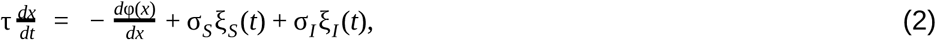

where the potential φ(*x*) = − μ*x* (Figure 1d-f, inset). The conceptual advantage of using a potential relies on the fact that the dynamics of the decision variable always “roll downward” towards the minima of the potential with only the fluctuations terms ξ_*S*_ (*t*) or ξ_*I*_ (*t*) causing possible motion upward. Notice that, although the term ξ_*S*_ (*t*) is modeled as a noise term, it represents the temporal variations of the stimulus which are under the control of the experimenter. The existence of decision bounds can be readily introduced in the shape of the potential, which strongly influences how stimulus fluctuations impact the upcoming decision: (1) in the DDMA (Figure 1a), absorbing bounds are implemented as two vertical “cliffs” such that when the decision variable arrives at one of them, it remains there for the rest of the trial. When this happens, the fluctuations late in the stimulus are unlikely to affect the decision, yielding a decaying psychophysical kernel (PK) characteristic of a “primacy” effect ^9,19,31,33,36^. (2) In the Drift Diffusion Model with Reflecting bounds (DDMR) (Figure 1b), the bounds are two vertical walls that set limits to the accumulated evidence; early stimulus fluctuations are largely forgotten once the decision variable bounces against one bound and hence the PK shows a “recency” effect ^19^. (3) In the Perfect Integrator (PI) (Figure 1c), there are no bounds, the stimulus is integrated uniformly across time yielding a flat PK ^18^. Thus, each of these three *canonical* models performs a qualitatively distinct integration process by virtue of how the bounds are imposed. Moreover, the characteristic integration dynamics of each model is invariant to changes in the stimulus parameters. To illustrate this, we show how the PKs depended on the magnitude of the stimulus fluctuations (*σ*_*S*_) (Figure 1). For very weak stimulus fluctuations, all three models are trivially equivalent because the bounds are never reached and hence the PKs are flat (Figure 1d-f). As *σ*_*S*_ increases, in both the DDMA and the DDMR, the bounds are reached faster yielding an increase and a decrease of the PK slope, respectively (Figure 1h). In these two models, the integration of evidence becomes more and more transient as *σ*_*S*_ increases, ultimately causing a decrease of the PK area (Figure 1g). The PK for the perfect integrator remains flat for all *σ*_*S*_ (zero PK slope, Figure 1h) and its area increases monotonically (Figure 1g). Thus, the dynamics of evidence accumulation are an invariant and distinct property of each model.

**Figure 1.**
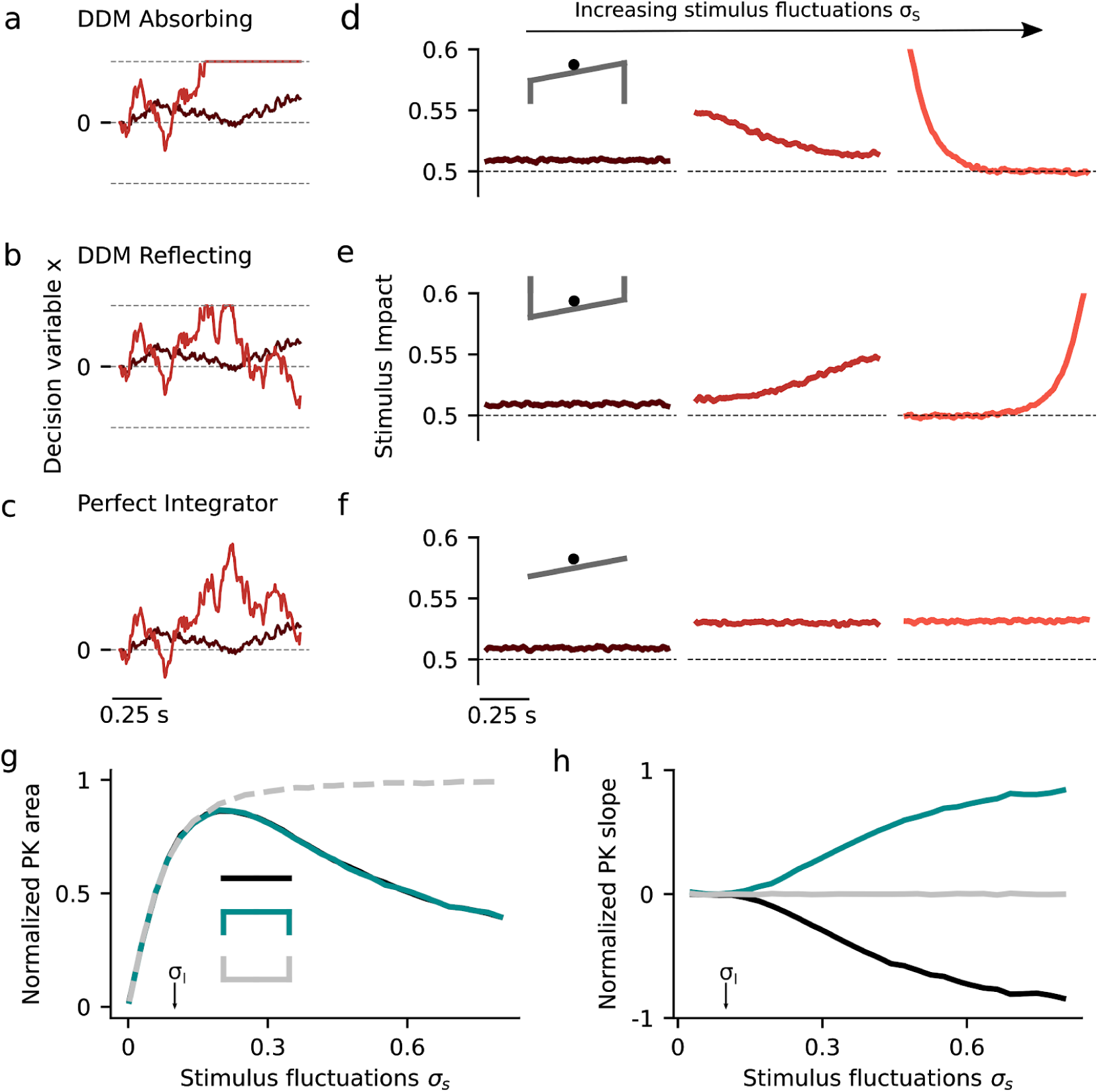
Dynamics of evidence accumulation in the canonical drift diffusion models. (**a-c**) Single-trial example traces of the decision variable *x(t)* for weak (*σ*_*S*_=0.09) and intermediate (*σ*_*S*_=0.25) stimulus fluctuations in the three canonical models. **a:** The DDM with absorbing bounds integrates the stimulus until it reaches one of the absorbing bounds represented in the potential landscape as infinitely deep attractors (see inset in d). The slope of the potential landscape is the mean stimulus strength, in this case (μ<0) **b**: The DDM with reflecting bounds integrates the stimulus linearly until a bound is reached when no more evidence can be accumulated in favor of the corresponding choice option (see inset in f). **c**: The perfect integrator integrates the entire stimulus uniformly, corresponding to a diffusion process with a flat potential (see inset in f). In the three models, the choice is given by the sign of *x(t)* at stimulus offset. (**d-f**) Psychophysical Kernels (PK) for the three canonical models for increasing magnitude of the stimulus fluctuations (from left to right): *σ*_*S*_=0.09, 0.25 and 0.53. (**g-h**) Normalized PK area and normalized PK slope as a function of *σ*_*S*_ for the three canonical models (see inset in g for color code). The area is normalized by the PK area of the perfect integrator with no internal noise (*σ*_*i*_=0) and hence measures the ability of each model to integrate the stimulus fluctuations. In all panels, internal noise was fixed at *σ*_*I*_=0.1 (see arrows in g and h) which was sufficiently small to prevent *x(t)* from reaching the bounds in the absence of a stimulus. Mean stimulus evidence was *μ*=0 in all cases.

### Neurobiological models show a variety of integration regimes

We next characterized the dynamics of evidence accumulation in the double well model (DWM), which can accurately describe the dynamics of a biophysical attractor network model ^20,28^. The DWM exhibits winner-take-all attractor dynamics defined by the non-linear potential *φ(x)*: 

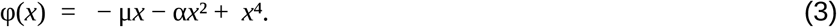

The resulting energy landscape can exhibit two minima (i.e. attractor states) corresponding to the two possible choices (Figure 2a, inset). The three terms of the potential, from left to right, capture (1) the impact of the net stimulus evidence *μ* which, as in the canonical models, tilts the potential towards the attractor associated with the correct choice; (2) the model’s internal categorization dynamics parameterized by the height of the barrier separating the two attractors (which scales with *α*^2^) and (3) bounds, also arising from the internal dynamics, that limit the range over which evidence is accumulated. We found that the DWM had a much richer dynamical repertoire as a function of stimulus fluctuations magnitude than the canonical models. Specifically, the attractors imposed the categorization dynamics but these could be overcome by sufficiently strong stimulus fluctuations. Thus, for weak σ_S_, the categorization dynamics dominated: when the system reached an attractor, it remained in this initial categorization until the end of the stimulus. In this regime, only early stimulus fluctuations occurring before reaching an attractor could influence the final choice, yielding a primacy PK (Figure 2c, light brown trace) (Wimmer et al. 2015; Wang 2002). For strong *σ*_*S*_, the initial categorization had no impact on the final choice because transitions between the attractors occurred readily. It was the fluctuations coming late in the trial which determined the final state of the system and hence the PK showed recency (Figure 2c, orange). For moderate values of *σ*_*S*_, there was an intermediate regime in which the PK was a mixture between primacy and recency, but not necessarily flat (Figure 2c, red line). We called this regime *flexible categorization* because it represented a balance between the internal categorization dynamics and the ability of the stimulus fluctuations to overcome their attraction (Figure 2b). As a result of this balance, the stimulus fluctuations impacted the choice over the whole trial (PK slope= 0; Figure 2e) because both initial fluctuations and later fluctuations causing transitions had a substantial impact on choice. Moreover, these fluctuations causing transitions were more temporally extended than those in the recency regime (Supplementary Figure 1a). Thus, the area of the PK reached its maximum (maximum area= 0.82; Figure 2f) implying that the integration of the stimulus fluctuations carried out by DWM was comparable to a perfect integrator (which has PK area equal 1). The same crossover from primacy to recency regimes, passing through the flexible categorization regime, could be achieved, at fixed *σ*_*S*_, by varying the stimulus duration (Figure 2d, g). This occurs because for a fixed magnitude of stimulus fluctuations, the *rate* of transitions was constant but the probability to observe a transition increased with the stimulus duration changing the shape of the PK accordingly (Figure 2d). In sum, depending on the capacity of the stimulus to generate transitions between attractors, the DWM model could operate in the primacy, the recency or the flexible categorization integration regime.

**Figure 2.**
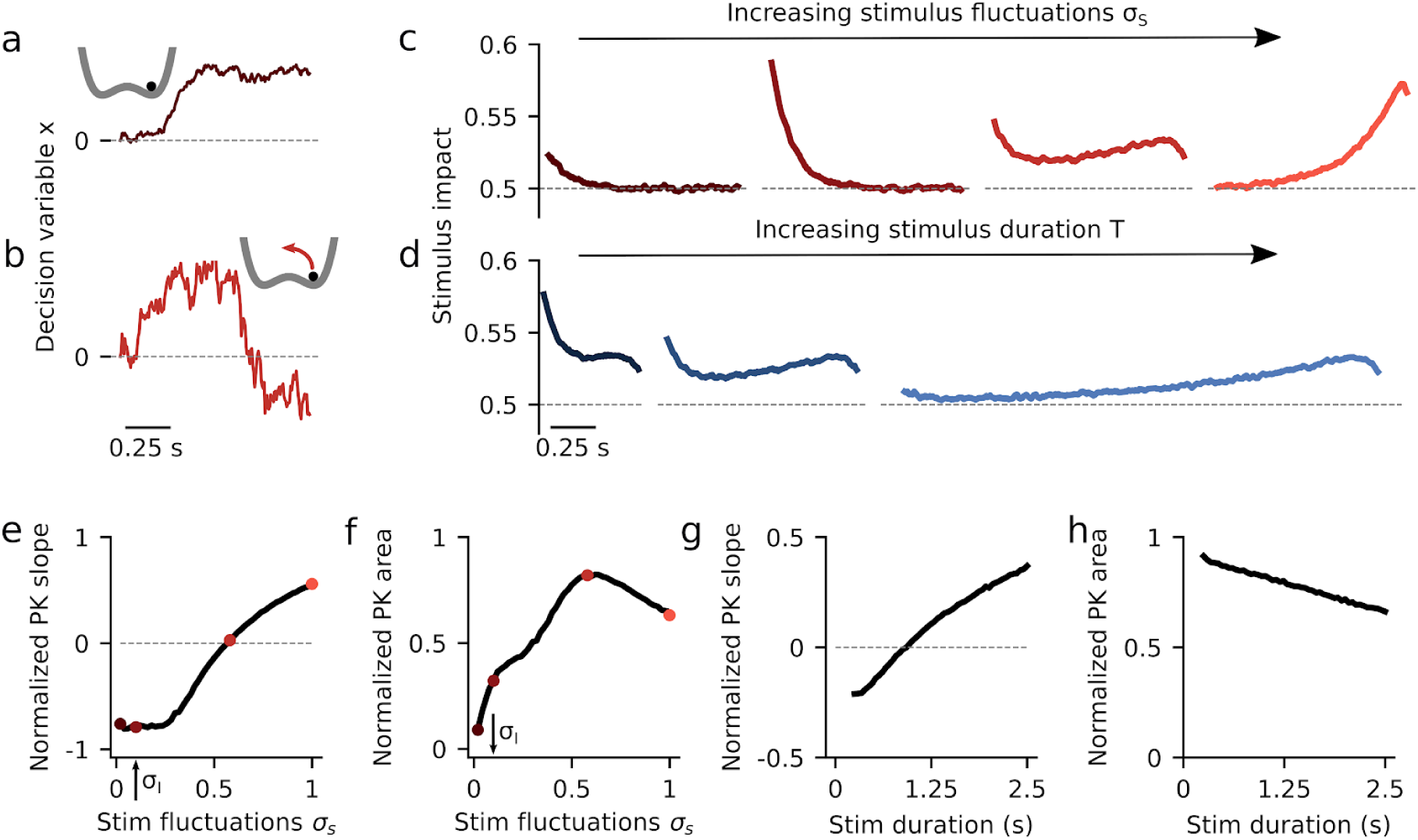
Dynamics of evidence accumulation in the double well model. (**a-b**) Single-trial example traces of the decision variable for the DWM with weak (a, *σ*_*S*_ = 0.1) and intermediate (b, *σ*_*S*_ = 0.58) stimulus fluctuations *σ*_*S*_. Transitions between attractors were only possible for sufficiently strong *σ*_*S*_ (insets). (**c**) PKs for increasing values of *σ*_*S*_ (from left to right, σ_S_= 0.02, 0.1, 0.58 and 1). (**d**) PKs for increasing values of stimulus duration *T* (from left to right, *T*= 0.5, 1 and 2.5 with σ_S_= 0.58). (**e-f**) Normalized PK slope and PK area as a function of σ_S_. Colored dots indicate the examples shown in panel c. The area peaks at the flexible categorization and it vanishes for small σ_S_ because choice is then driven by internal noise. (**g-h**) Normalized PK slope and area as a function of *T* with σ_S_= 0.58. As *T* increases, the DWM integrates a smaller fraction of the stimulus making the area decrease monotonically. Internal noise was σ_*I*_ = 0.1 in all panels (see arrows in panels e and f).

### Decision accuracy in models of evidence integration

Given that the DWM changes its integration regime when *σ*_*S*_ is varied, we next investigated the impact of this manipulation on the decision accuracy. We set the internal noise to *σ*_*I*_= 0 and computed the psychometric function *P*(*μ*,*σ*_*S*_) showing the proportion of correct choices as a function of the mean stimulus evidence μ and the strength of stimulus fluctuations *σ*_*S*._ For small fixed *σ*_*S*_ the section of this surface yielded a classic sigmoid-like psychometric curve *P*(*μ*) (Figure 3a, dark brown curve). As *σ*_*S*_ increased, this curve became shallower simply because larger fluctuations implied a drop in the signal to noise ratio of the stimulus (Figure 3a, red and orange curves). Unexpectedly, however, the decline in sensitivity of the curve *P*(*μ*) was non-monotonic (Figure 3a), an effect which was best illustrated by plotting the less conventional psychometric curve *P*(*σ*_*S*_) at fixed *μ* (Figure 3a-b, black curve). To understand this non-monotonic dependence, we first defined two transition probabilities: the *correcting* transition probability *p*_*C*_ was the probability to be in the correct attractor at the end of a trial, given that the first visited attractor was the error. The *error-generating* transition probability *p*_*E*_ was the opposite, i.e. the probability to finish in the wrong attractor given that the correct one was visited first (see Methods). Using Kramers’ reaction-rate theory ^37^ the transition probabilities could be analytically computed, and the accuracy *P* could be expressed as the probability to initially make a correct categorization and maintain it, plus the probability to make an initial error and reverse it:

**Figure 3.**
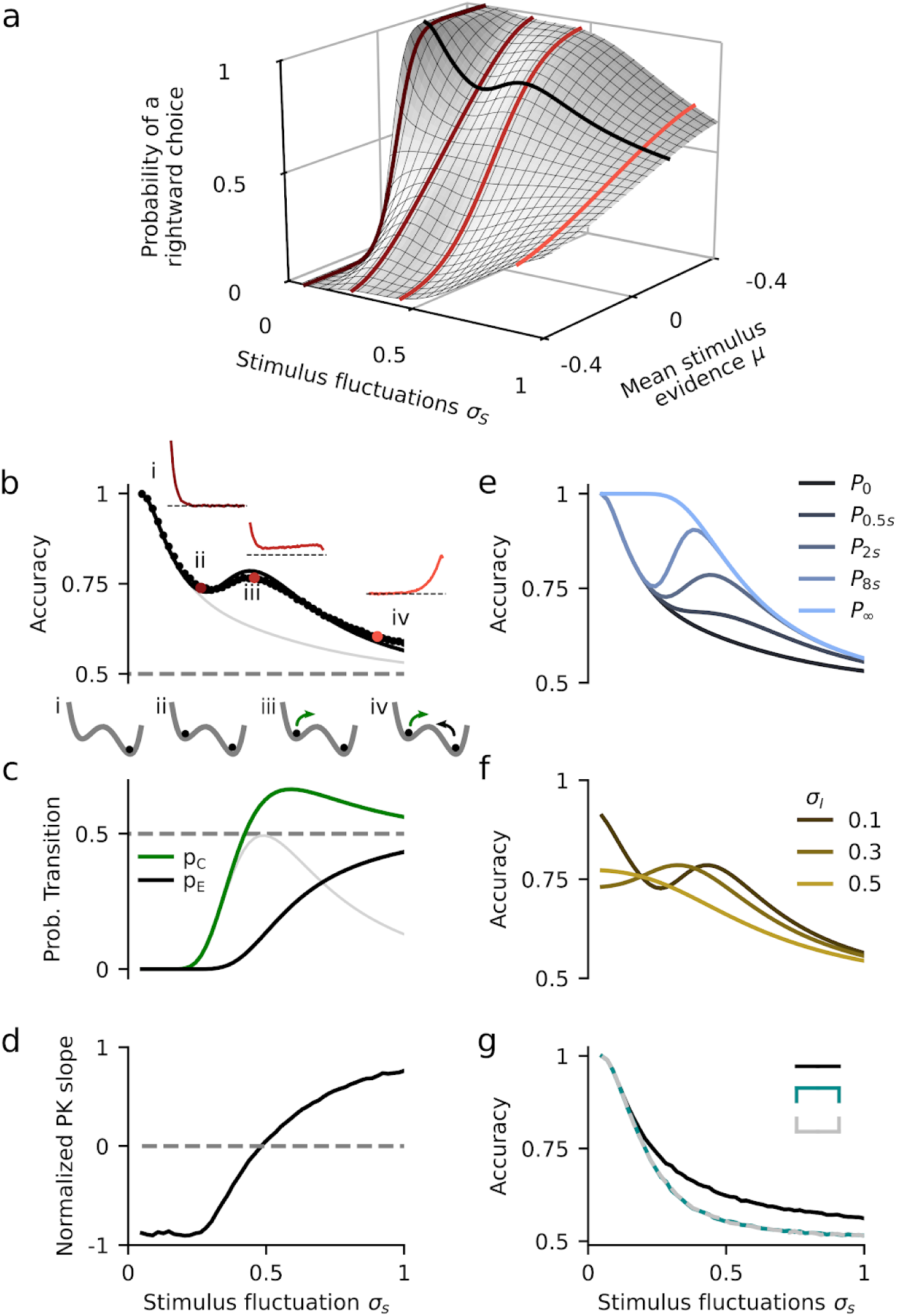
Impact of stimulus fluctuations on choice accuracy in the double well model. (**a**) Probability of a righward choice as a function of the mean stimulus evidence (*μ*) and the stimulus fluctuations (*σ*_*S*_). The colored lines show *classic* psychometric curves, accuracy vs. *μ* (for fixed *σ*_*S*_= 0.07, 0.26, 0.46 and 0.90) whereas the black line shows the accuracy vs. *σ*_*S*_ (for fixed *μ*= 0.15). (**b**) Accuracy (*P*) as a function of the stimulus fluctuations *σ*_*S*_ obtained from numerical simulations (dots) and theory (line, same as black line in **a**). Insets show the PK for three values of *σ*_*S*_ (marked with colored dots). The grey line shows the accuracy of the first visit attractor (*P*_*0*_). (**c**) Probability to make a correcting *p*_*C*_ (green) or an error transition *p*_*E*_ (black) and their difference *p*_*C*_ -*p*_*E*_ (gray). The local maximum in *P* coincides with the maximum difference between the two probabilities. Insets: sequence of regimes as transitions become more likely: i) For negligible *σ*_*S*_, the decision variable always evolves towards the correct attractor; ii) as *σ*_*s*_ increases, the decision variable can visit the incorrect attractor but neither kind of transition is activated; iii) for stronger *σ*_*S*_, only the correcting transitions (green arrow) are activated; iv) for strong *σ*_*S*_, both types of transition are activated. (**d**) Normalized PK slope as a function of *σ*_*S*_. The flexible categorization regime, reached when the index is close to zero, coincides with the local maximum in accuracy (a). (**e**) Accuracy versus *σ*_*S*_ for different stimulus durations *T* (see inset). The accuracy for any finite *T* shifts as *σ*_*S*_ increases between the probability to first visit the correct attractor *P*_*0*_ and the stationary accuracy *P*_*∞*_. (**f**) Accuracy versus *σ*_*S*_ for different magnitudes of the internal noise (see inset). (**g**) Accuracy versus *σ*_*S*_ for the three canonical models (see inset). The internal noise was σ_*i*_ = 0 in all panels except in f.

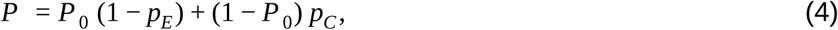

where *P*_*0*_ was the probability of first visiting the correct attractor (Methods). When the fluctuations were negligible *σ*_*S*_≈0, the decision variable always rolled down towards the correct choice because the double well potential was tilted to the correct attractor (e.g. *μ*>0), and hence *P* = 1 (Figure 3b *i*). As *σ*_*S*_ started to increase, fluctuations early in the stimulus could cause the system to fall into the incorrect attractor but, because fluctuations were not sufficiently strong to generate transitions (*p*_*E*_ ≈ *p*_*C*_ ≈ 0), accuracy was *P=P*_*0*_ (Equation 4) and decreased with *σ*_*S*_ towards chance (gray line in Figure 3b). As the stimulus fluctuations grew stronger, the transitions between attractors became more likely but, because the barrier to escape from the incorrect attractor was smaller than the barrier to escape from the correct attractor, the two transition probabilities were very different. Specifically, Kramers’ theory shows that the ratio between *p*_*C*_ and *p*_*E*_ depends exponentially on the barrier height difference (see Methods). Thus, *p*_*C*_ increased steeply with *σ*_*S*_, even before *p*_*E*_ reached non-negligible values (Figure 3c) opening a window in which transitions were *only* correcting: accuracy became *P* ≃ *P* _0_ + (1 − *P* _0_) *p*_*C*_ and it increased with *σ*_*S*_ (Figure 3b *iii*). The maximum difference between *p*_*C*_ and *p*_*E*_ coincided with the flexible categorization regime in which the PK slope was zero and the accuracy showed a local maximum (Figure 3b-d). Finally, for strong *σ*_*S*_, error transitions also became likely and the net effect of stimulus fluctuations was again deleterious, causing a decrease of *P*. In sum, it was the large difference in transition probabilities caused by the double well landscape which led to the non-monotonic dependence of the psychometric curve. Because the canonical models lacked attractor dynamics, the accuracy in all of them decayed monotonically with the stimulus fluctuations (Figure 3g).

We next asked whether the non-monotonicity of the psychometric curve was robust to variation of other parameters such as the mean stimulus evidence *μ*, the stimulus duration *T* and the internal noise *σ*_*I*_. We found that the non-monotonicity was robustly obtained over a broad range of *μ*, ranging from small values just above zero to a critical value beyond which the curve became monotonically decreasing (Supplementary Figure S2). Because the transition probabilities scale with the stimulus duration *T*, the psychometric curve *P(σ*_*S*_) was strongly affected by changes in *T* (Figure 3e). To understand this dependence, we rewrote the transitions probabilities *p*_*C*_ and *p*_*E*_ from Equation 4 as a function of the transition rates and the stimulus duration (see Methods, Equations 17 and 18): 

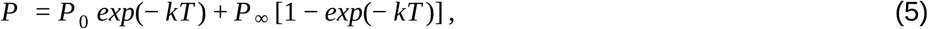

where *k* is the sum of the transition rates from both attractors and *P*_*∞*_ is the stationary accuracy (i.e. the limit of *P* when *T* → ∞). As expected, the two psychometric curves *P*_*0*_ and *P*_*∞*_, which decreased monotonically with *σ*_*S*_, delimited the region in which *P* existed: for weak *σ*_*S*_, *P* followed the decay of the psychometric curve *P*_*0*_, whereas for strong *σ*_*S*_ it tracked the decay of the stationary accuracy *P*_*∞*_. The switching point occured when the probability to observe a transition was substantial, i.e. when *kT* ∼ 1. For longer stimulus durations, the activation of the transitions occurred for weaker *σ*_*S*_ and consequently the bump in accuracy was shifted towards the left also becoming more prominent (Figure 3e, Methods). For very short *T*, the activation of the transitions occurred for such a large value of *σ*_*S*_ that the two curves *P*_*0*_ and *P*_*∞*_ have come too close and the psychometric *P(σ*_*S*_) was then monotonically decreasing (Figure 3e). Finally, when we set the internal noise to a non-zero value, it sets a minimal level of fluctuations below which no stimulus magnitude *σ*_*S*_ could go, effectively cropping the psychometric curve *P(σ*_*S*_) from the left (Figure 3f). Only when the internal noise was larger than a critical value the psychometric curve became monotonically decreasing (Supplementary Figure S2, see Methods for the computation of the critical noise value). In sum, the non-monotonicity of the psychometric curve was a robust effect, being most prominent for values of the mean stimulus evidence *μ* yielding an intermediate accuracy (i.e. *P* ∼ 0.75), long stimulus durations and weak internal noise.

### Consistency in models of evidence integration

In order to identify further signatures of the nonlinear attractor dynamics that could be tested experimentally, we studied the choice consistency of the double well model (DWM). Choice consistency is defined as the probability that two presentations of the same exact stimulus, i.e. the same realization of the stimulus fluctuations, yield the same choice. In the absence of internal noise, the decision process in the model is deterministic and consistency is 1. In contrast, when the stimulus has no impact on the choice, the consistency is 0.5. We used the double-pass method, which presents each stimulus twice ^13,38,39^, to explore how consistency in the DWM depended on *σ*_*S*_ and *σ*_*I*_ (Figure 4). We only used *μ*=0 stimuli with exactly zero integrated evidence in order to avoid the parsimonious increase of consistency due to larger deviations of the accumulated evidence from the mean (see Methods). As expected, consistency was close to 0.5 when *σ*_*S*_ was small compared to *σ*_*I*_, and it increased with increasing *σ*_S_ (Figure 4a). However, despite this general increase, we found a striking drop in consistency for a range of intermediate *σ*_*S*_ values. Thus, consistency could depend non-monotonically on the strength of stimulus fluctuations, a similar effect as observed for choice accuracy. To understand this effect, we studied the time-course of the decision variable *x* over many repetitions of a single stimulus, at different values of *σ*_*S*_ (Figure 4d-h). For very small *σ*_*S*_, consistency was 0.5 because the internal noise was the dominant factor making both choices equally likely (Figure 4d). As *σ*_*S*_ grew, stimulus fluctuations could determine the first visited attractor but decision reversals were still not activated, yielding a high consistency (Figure 4e). For larger *σ*_*S*_, transitions occurred but only when internal noise and the stimulus fluctuations worked together to produce a large fluctuation (Figure 4f). The necessary contribution of the internal noise, that varied from trial to trial, led to the decrease in consistency. Once *σ*_*S*_ was large enough to cause reversals on its own, consistency increased again (Figure 4g). Thus, as with the non-monotonicity in the psychometric curve, it was the difference between two transition probabilities, the transition probability with internal noise versus the probability without internal noise, that was maximal when consistency decreased (Figure 4b). Also as before, to observe the non-monotonicity in the consistency, *σ*_*I*_ had to be sufficiently small not to cause transitions on its own (Figure 4a-b). Notice however that the non-monotonicity here was not caused by the asymmetry between correcting versus error transitions, as consistency was computed using μ=0 stimuli (i.e. there was no correct choice). The effect was a result of the nonlinear attractor dynamics of the DWM and thus it could not occur in any of the canonical models (Figure 4c).

**Figure 4.**
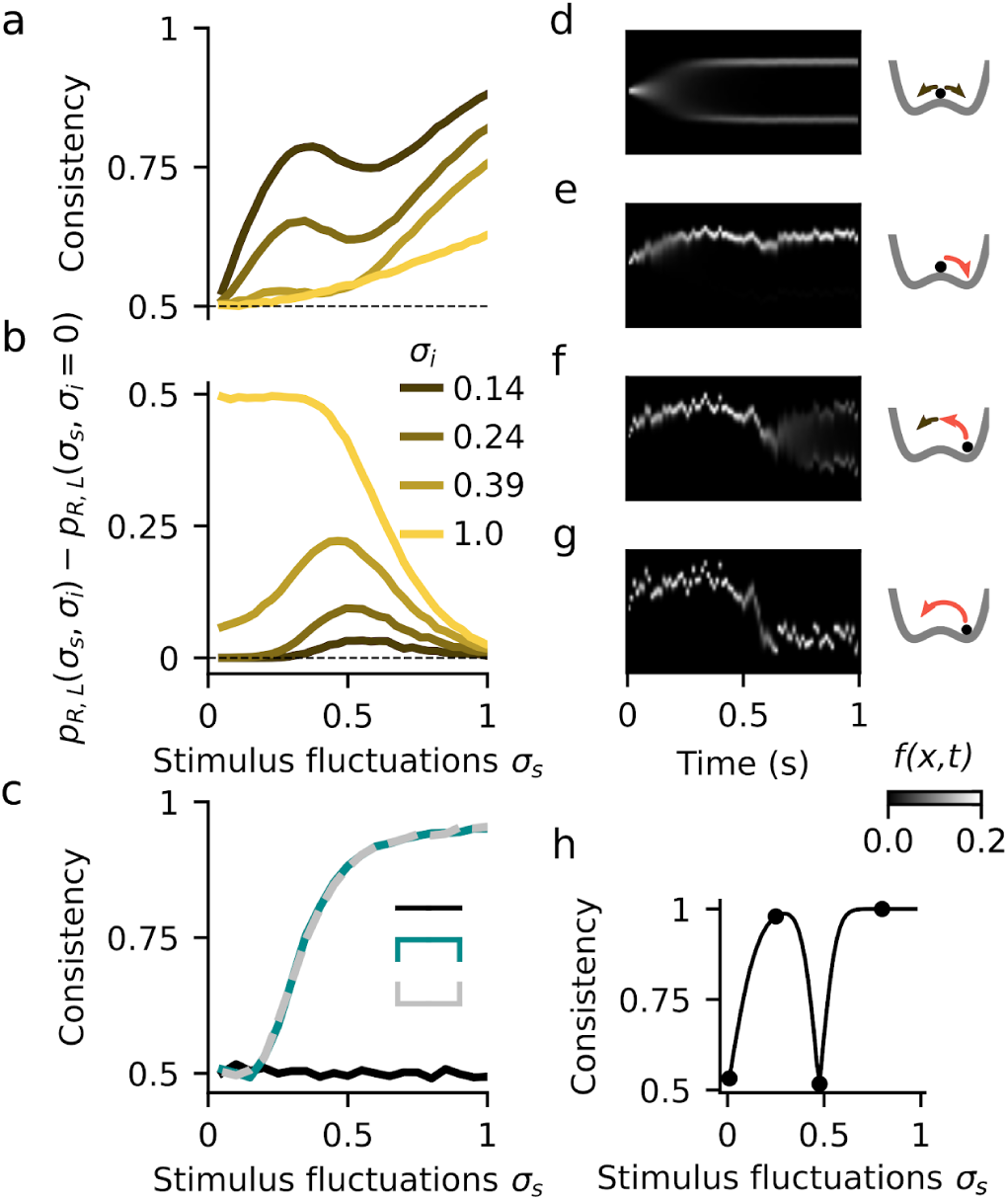
Dependence of choice consistency on stimulus fluctuations. (**a**) Average consistency versus stimulus fluctuations *σ*_*S*_ for different values of the internal noise *σ*_*I*_ (see inset in b). (**b**) Difference between the transition probabilities with (*p*_*R*,*L*_(*σ*_*S*_,*σ*_*I*_)) and without ((*p*_*R*,*L*_(*σ*_*S*_,*σ*_*I*_=0))) internal noise. The drop in consistency coincides with an increase of this difference revealing the *σ*_*S*_-range in which transitions occurred because of the cooperation of internal and stimulus fluctuations. **(c)** Consistency versus *σ*_*S*_ for the canonical models. The consistency of the perfect integration is at chance level because we used stimuli with exactly zero integrated evidence (see Methods). (**d-g**) Temporal evolution of the decision variable probability distribution *f(x*,*t)* for an example stimulus in the different regimes of *σ*_*S*_: for negligible *σ*_*S*_ the choice is driven by the internal noise and the consistency is very low (53.2%, d). For small *σ*_*S*_, when the stimulus determines the first visited attractor but fluctuations are not strong enough to produce transitions, the consistency is very high (97.8%, e) For intermediate *σ*_*S*_, the transitions can only occur when *σ*_*I*_ and *σ*_*S*_ work together to cause a large fluctuation. Because the internal noise has again impact on the choice, the consistency decreases (65.5%, f). For large *σ*_*S*_, the stimulus fluctuations are strong enough to produce transitions by itself and the consistency is again very high (100%, g) (**h**) Consistency vs. *σ*_*S*_ obtained just using the example stimulus shown in d-g (points mark the *σ*_*S*_ values shown in d-g). Mean stimulus evidence was *μ* =0 in all panels.

### Flexible categorization in a spiking network with attractor dynamics

Having shown that the DWM generates signatures of attractor dynamics which are qualitatively different from any canonical model, we then assessed whether these could be reproduced in a more biophysically realistic network model composed of leaky integrate-and-fire neurons (Methods). The network consisted of two populations of excitatory (E) neurons (N_E_= 1000 for each population), each of them selective to the evidence supporting one of the two possible choices, and a nonselective inhibitory population (N_I_= 500) (Figure 5a). The network had sparse, random connectivity within each population (probability of connection between neurons was 0.1). The stimulus was modeled as two fluctuating currents, reflecting evidence for each of the two choice options and injected into the corresponding E population. The two currents were parametrized by their mean difference *μ* and their standard deviation *σ*_*S*_ (see Methods). In addition, all neurons in the network received independent stochastic synaptic inputs from an external population. As in previous attractor network models used for stimulus categorization, the two E populations competed through the inhibitory population ^20^. Thus, upon presentation of an external stimulus, there were two stable solutions: one in which one E population fired at a high rate while the other fired at a low rate and vice versa (Figure 5). Similar to the DWM, we found a non-monotonic relation between the accuracy and the magnitude of the stimulus fluctuations *σ*_*S*_ provided the stimulus duration *T* was sufficiently long (Figure 5b). Moreover, as *σ*_*S*_ increased the integration regimes of the network changed from primacy to recency, passing through the flexible categorization regime (Figure 5c-f). In this regime, transitions between attractor states occurred when there were input fluctuations that extended over hundreds of milliseconds, indicating that the temporal integration of evidence continued even after one of the attractors was reached (Supplementary Figure S1b). The crossover between primacy and recency regimes was also observed at constant *σ*_*S*_ when we varied the stimulus duration *T* (Figure 5g). Thus, the signatures of attractor dynamics that we found in the DWM were replicated in an attractor network with biophysically plausible parameters.

**Figure 5.**
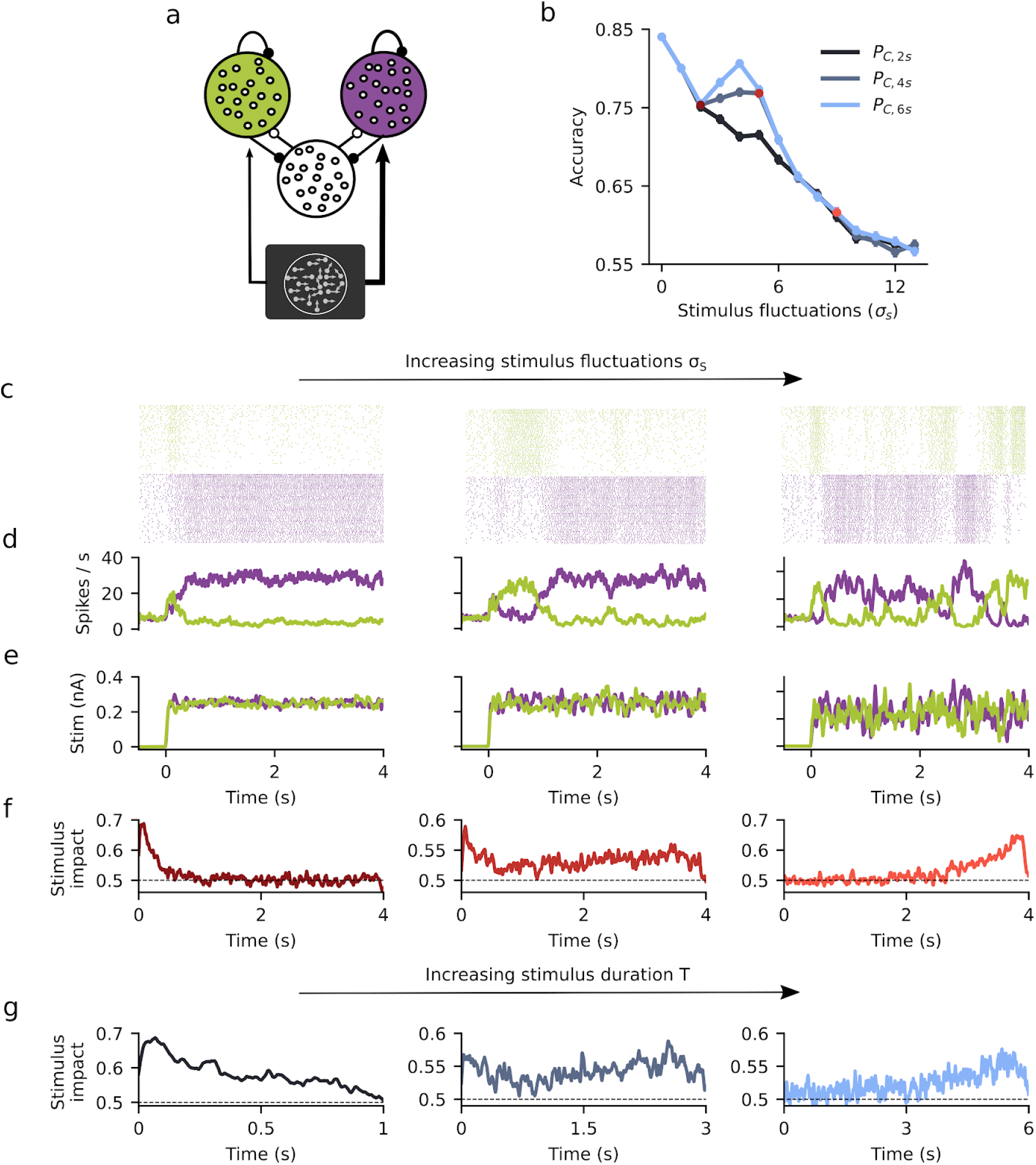
Signatures of winner-take-all attractor dynamics in a spiking network. (**a**) Schematic of the spiking network consisting of two stimulus-selective populations (green and purple) made of excitatory neurons that compete through an untuned inhibitory population (white population). (**b**) Accuracy *P*_*C*_ versus stimulus fluctuations σ_*S*_ obtained from simulations of the spiking network for three values of the stimulus duration *T* =2, 4 and 6 seconds (see inset). (**c-e**) Single trial examples showing spike raster-gram from the two excitatory populations (c), traces of the instantaneous population rates (d) and of the input stimuli (e), for different values of stimulus fluctuations σ_*S*_ = 2 (left), 5 (middle) and 9 pA (right). Colored points in (b) indicate the σ_*S*_ used. **(f)** Psychophysical kernels obtained for each σ_*S*_ value. The mean input difference was μ = 0.03 pA and the stimulus duration *T* = 4 s. (**g**) Psychophysical kernels for different stimulus duration *T* = 1, 3 and 5 s, from left to right.

### Changes in PK with stimulus duration in human subjects unveiled the flexible categorization regime

We tested whether the DWM could parsimoniously account for the variations of the integration dynamics previously found in a perceptual categorization task as the stimulus duration was varied ^34^. In the experiment, human subjects had to discriminate the brightness of visual stimuli of variable duration *T* = 1, 2, 3 or 5 s. Confirming previous analyses ^34^, the average PKs across subjects changed from primacy to recency with increasing stimulus durations (Figure 6a). To assess whether these changes in the shape of the PKs could be captured by the DWM, we used the DWM to categorize the same stimuli (the exact same temporal stimulus fluctuations and number of trials; see Methods) that were presented to the human subjects (Figure 6c-f). We found that the PKs for different stimulus durations obtained in the DWM were very similar to the experimental data (Figure 6b). Importantly, these results were obtained with fixed model parameters for all stimulus durations suggesting that the variation in PK did not necessarily indicate a change of the integration mechanism of the model, as previously suggested ^34^. Rather, fixed, but nonlinear attractor dynamics in the DWM parsimoniously accounted for the observed PK changes.

**Figure 6.**
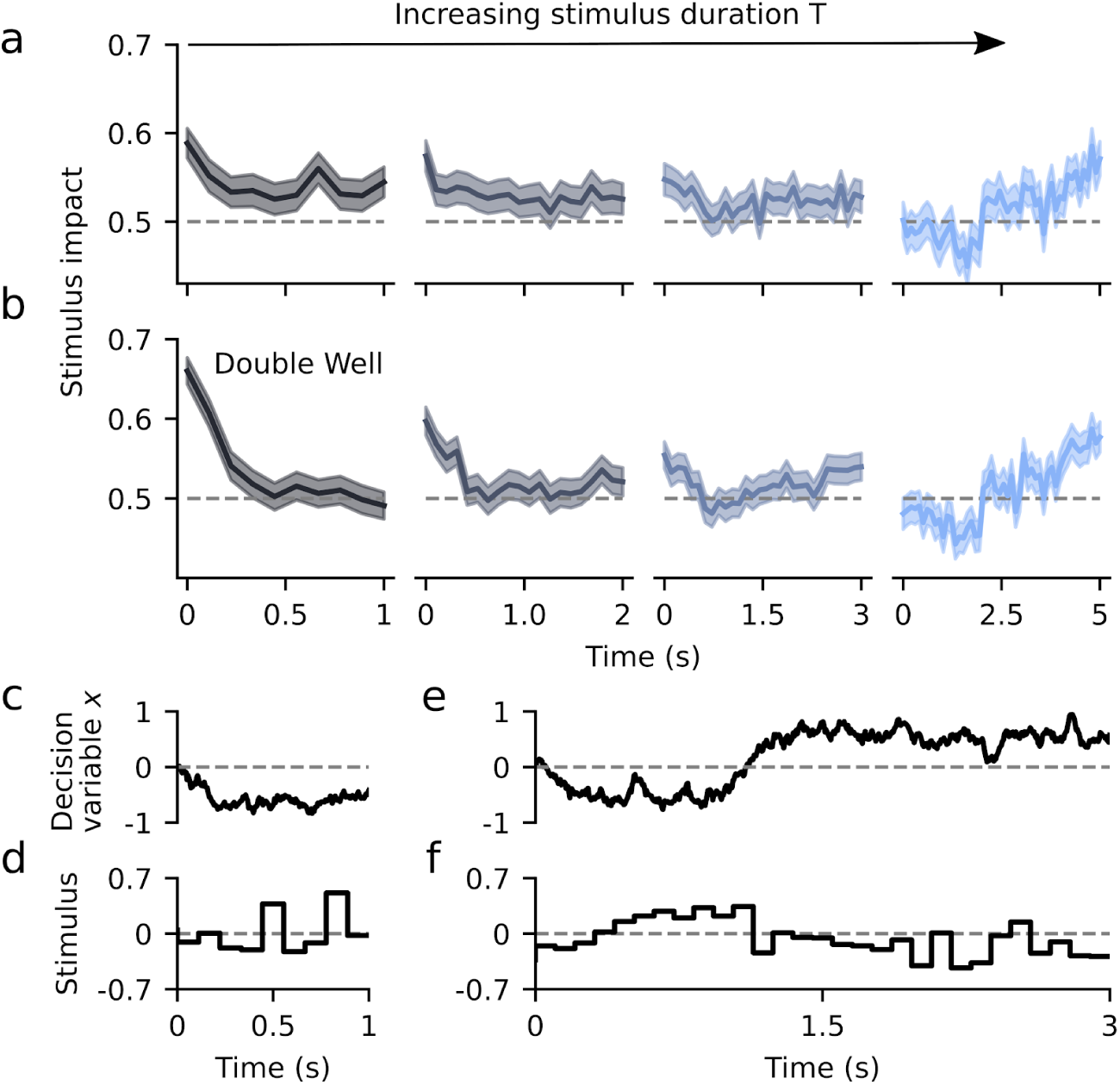
The double well model accounts for experimentally observed changes in psychophysical kernels. **(a)** Psychophysical kernels for different stimulus durations, obtained from human subjects performing a brightness discrimination task (N= 21) ^34^. From left to right, stimulus duration was *T*= 1, 2, 3 and 5 seconds. **(b)** Psychophysical kernels obtained by fitting the DWM to categorize the very same stimuli presented to the human subjects (i.e. same temporal fluctuations of net evidence; see Methods). Lines represent the kernels obtained from pooling all data across subjects and the error bands represent s.e.m. **(c-f)** Example traces of the decision variable of the fitted DWM (c,e) and the stimulus (d,f) for 1 and 3 s trials.

### Stimulus integration across a memory period is consistent with flexible categorization dynamics

Finally, we tested the DWM in a task that requires evidence accumulation and working memory. We used published data from two studies carrying out a psychophysical experiment in which subjects had to categorize the motion direction of a random dot kinematogram ^35,40^.

Interleaved with the trials showing a single kinematogram (single pulse trials, duration 120 ms) there were also trials having two kinematograms separated by a temporal delay (two pulse trials). In these two pulse trials, subjects had to combine information from both pulses in order to categorize the average motion direction. The two pulses could have different motion coherence but they always had the same motion direction. Subjects were able to combine the evidence from the two pulses and their accuracy did not depend on the duration of the delay period for durations up to 1 s, meaning that they were able to maintain the evidence from the first pulse without memory loss. Overall, subjects gave slightly more weight to the second than the first pulse (Primacy-Recency Index=0.22; see Methods). Qualitatively, the DWM could in principle capture this behavior because its underlying dynamics can solve the two parts of the task, the maintenance of information during the working memory period and the combination of the two pulses of evidence (Figure 7a). The model would categorize the first pulse in one of the attractors, which would be stably maintained during the delay because the internal noise is insufficient to cause transitions.

**Figure 7.**
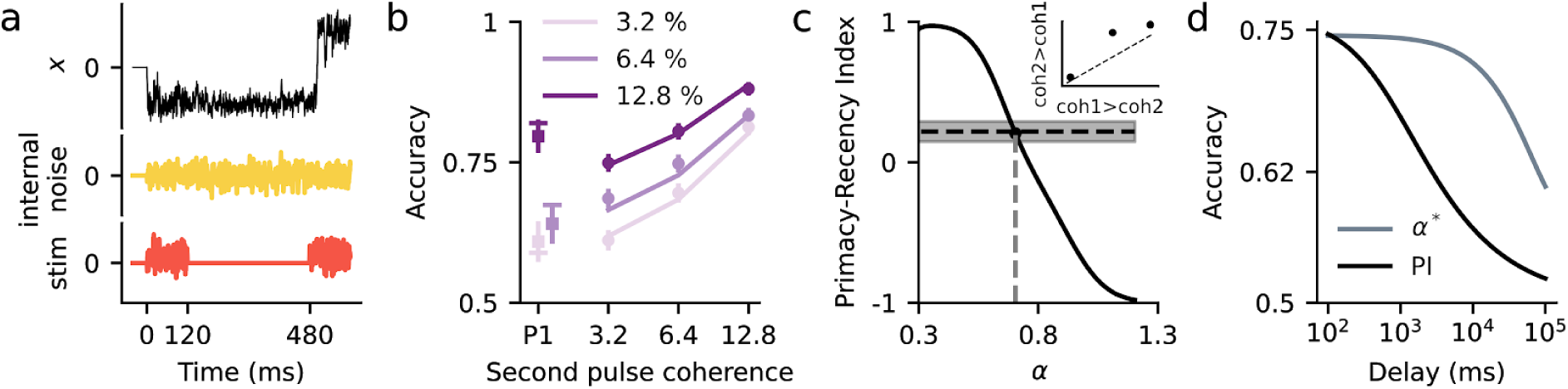
The flexible categorization regime accounts for the combination of two pulses of evidence during a working memory task. (**a**) Traces of the decision variable of the DWM (black), the internal fluctuations (yellow) and the stimulus (orange) for an example double pulse trial. (**b**) Accuracy for single (squares) and two pulse trials (dots) versus the coherence of the second pulse observed in the data from ^35,40^ (dots) and the values obtained from the fitted DWM (lines). Because accuracy in the experiment did not depend on delay length, dots show the average accuracy across all delays. Different colors represent different first pulse coherences (see inset). Symbols show mean across subjects and error bars show 95% confidence intervals. (**c**) Primacy-recency index (PRI) for the DWM as a function of the barrier height (*c*_2_). The black dot marks the PRI for the fitted parameter *α\**= 0.7. The horizontal line is the PRI computed from the psychophysical data (grey area 95% confidence interval). Inset: accuracy for two pulse stimuli in which coherence is larger in pulse 2 than in pulse 1 (i.e. coh2 > coh1) versus accuracy for the same pulses presented in the reverse order (i.e. coh1>coh2). Consistent with the recency effect, accuracy is slightly better for coh2>coh1 stimuli. (**d**) Accuracy as a function of the delay duration for DWM and for the Perfect Integrator. In the DWM, which used the fitted parameter *α\** and σ_*i*_ = 0.32 and σ_*S*_ = 0.40, the accuracy is independent of the delay up to 1 s. In contrast, for the same internal noise σ_*i*_, the accuracy of the perfect integrator decreases continuously for all delays.

Finally, given the asymmetry in the DWM transition rates (Figure 3c), the second pulse could reverse incorrect initial categorizations while minimizing the risk of erroneously reversing correct ones (Figure 7a). To assess whether the DWM could indeed fit the data quantitatively, we computed the accuracy for each stimulus condition using Kramers’ transition rate theory and fitted the parameters using maximum likelihood estimation (solid lines, Figure 7b; Methods). We found that the DWM could fit the accuracy across conditions quite accurately (Figure 7b). Interestingly, the fitted DWM worked close to the flexible categorization regime, matching the slight recency effect coming out from the combination of the two pulses (Figure 7c).

Because subjects’ accuracy did not depend on delay duration ^35,40^, the model fitting could only determine the value of the sum of the stimulus and internal noises 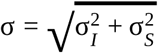 and set an upper bound 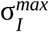 for the internal noise: for any value 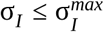 the transitions during the delay were negligible (< 1%) and the DWM yielded the same behavior (see Methods). Choosing the σ_*I*_ to be at the upper bound 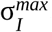, yielded a constant accuracy for delays up to ∼1 s. For longer delays, however, transitions during the delay became active causing forgetting and accuracy decrease (Figure 7d) as has been shown in experiments using a broader range of delays (Melcher et al. 2004). In contrast, the perfect integrator did not show a range of delays over which the accuracy remained constant (Figure 7d): the internal noise had a much larger impact on the maintenance of stimulus evidence so that, for any significant level of internal noise, the accuracy decreased continuously with delay duration. In total, our analysis shows that the DWM can quantitatively fit psychophysical data from a working memory task, and that longer delays could provide a qualitative test for the model.

## Discussion

We have investigated the attractor model with winner-take-all nonlinear dynamics and we have found new, experimentally testable signatures that can distinguish it from the other models. First, the attractor model exhibits a continuous crossover from the primacy regime ^20,24^ to the recency regime. Between these two regimes we found the new flexible categorization regime in which the integration of stimulus fluctuations was maximally extended over time (Figure 2; Supplementary Figure 1). Second, in this regime a qualitative asymmetry between correcting and error transitions gave rise to a non-monotonic psychometric curve (Figure 3). Third, the rapid activation of transitions between decision states with the stimulus fluctuations also caused an unexpected non-monotonic dependence of the stimulus consistency (Figure 4a). Finally, we used two previous psychophysical experiments to show that the attractor model can quantitatively fit variations in PK profile with stimulus duration (Figure 5) and fit categorization accuracy in a task with integration of evidence across memory periods (Figure 6).

Recently, two studies have proposed alternative models that can explain the differences of PK time-courses found across subjects and experiments. In the first model, based on approximate Bayesian inference, the primacy effect produced by bottom-up vs. top-down hierarchical dynamics, was modulated by the stimulus properties which could yield different PK time-courses, a prediction that was tested in a visual discrimination task ^41^. The second study proposed a model that can produce different PK time-courses by adjusting the time scales of a divisive normalization mechanism, which yields primacy, and a leak mechanism, which promotes recency ^42^. In addition, this model can also account for bump shaped PKs, a class of PK that was found together with primacy, recency and flat PKs, in a study carried out using a large cohort of subjects (>100) ^43^. In the attractor model, the differences in the PK found across subjects or fixed stimulus properties could be explained by individual differences in the shape of the potential. Specifically, differences in the height of the barrier between the two attractor states would generate a variety of PK time-courses (Figure 7c) as the integration regime ultimately depends on the ratio between the total noise 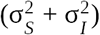 and the height of the barrier. A natural extension of our approach would be to assume that a time-varying process during the trial, e.g. an urgency signal ^44^, can progressively modify the shape of the potential. In that case, the DWM with an urgency signal ^44^ that changed the shape of the potential from a single well at stimulus onset into a double well at stimulus offset could readily reproduce the bump shaped PKs (not shown) recently reported ^43^. In sum, the attractor model shows a large versatility generating the diversity of PK shapes reported in the literature ^9,11,13,16–18,43^. Although several distinct models can account for the variety of PK shapes, they rely on a variety of neural mechanisms. Future electrophysiological or psychophysical experiments where the different models predict qualitatively different results will help distinguish between these possible mechanisms.

It has been previously shown that noise, from the stimulus or internal sources, can increase the accuracy of an attractor model with three stable attractors (i.e. with multistability): an undecided state and two decision states ^45,46^. In this model, the decision variable starts in the undecided state and, if it does not escape from this state during the stimulus presentation, the decision is made randomly. Thus, the noise can allow the decision variable to escape from the undecided state and increase the accuracy. Here, we have studied the attractor model in the winner-take-all regime, i.e. without an undecided state, and we have found that it is the large difference between the rate of correcting and error-generating transitions that produces the increase in accuracy in the flexible categorization regime. This is conceptually very different from transitions between the undecided state to the decision states. The same mechanism presented here drives the classic stochastic resonance ^47^ where a particle moving in a double well potential driven by a periodic signal necessitates of a suitable magnitude of noise for the system to follow the signal (i.e. escape from the well when it is no longer the global minimum). Similar to the effect described with the multistable attractor model ^45^, the accuracy decreases to chance in the deterministic noiseless case (σ = 0). In contrast, the accuracy for the DWM is greatest for σ = 0 because the initial position of the decision variable (*x*_0_ = 0) belongs to the basin of attraction of the correct attractor and thus it always rolls down to the correct attractor. However, whether this bump in accuracy produced by the attractor model as a function of the stimulus fluctuations (σ_*S*_) is a local or a global maximum, or if it exists at all, depends on internal parameters such as the internal noise (σ_*I*_) or the height of the barrier. These internal parameters can be different for different subjects and thus, one should expect to find this non-monotonic psychometric curve only in a fraction of subjects. Indeed, we carried out a visuospatial binary categorization task in which the fluctuations of the evidence σ_*S*_ were varied systematically from trial-to-trial. Preliminary analysis shows that the majority of subjects display a psychometric curve *P* (σ_*S*_) with a plateau followed by a decay as σ_*S*_ increased. A fraction of subjects exhibited however a non-monotonic dependence but the dependence of PK and other aspects of their behavior (e.g. idiosyncratic biases) on σ_*S*_ were not fully captured by the DWM dynamics. A future study will extend the DWM so that it can capture these data (G.P.O. manuscript in preparation).

The key mechanism underlying the flexible categorization regime are the transitions between attractor states which, functionally, can be viewed as changes of mind ^6^. Changes of mind have been previously inferred from sudden switches in the direction of the motor response ^6,48^ but also from decision bound crossings of the decision variable read out from neuronal population recordings ^49–52^. In reaction time tasks, an extension of the drift diffusion model can fit the modulation of the probability of observing a change of mind as a function of the mean stimulus strength ^6,48^. In this model, a first crossing of the decision bound initializes the response that is reversed if the decision variable crosses the opposite bound before the motor response is completed. As the DWM, this model predicts that correcting changes of mind are more likely than error changes of mind. However, this asymmetry does not imply a non-monotonic accuracy with the stimulus fluctuations in a fixed duration task. This is because, in the linear DDM with changes of mind ^6^, the correcting transition probability *p*_c_ is not exponentially more likely than error transitions as in the DWM (Equation 20). Thus, the benefit of having more correcting transitions as σ_*S*_ increases does not offset the cost of decreasing the signal-to-noise ratio (not shown). An attractor network has also been used previously to explain changes of mind during the motor response ^53^. Our work extends this study in several ways, by characterizing the full spectrum of integration regimes in the attractor model and by showing qualitative experimentally testable signatures of decision state transitions (e.g. non-monotonicity in the accuracy and coherence vs. σ_*S*_).

An important question in perceptual decision making is the extent to which subjects can integrate evidence during the stimulus presentation. It has been recently pointed out that differentiating between integrating and non integrating strategies may be more difficult than naively thought ^54^. Here we evaluate the degree of evidence integration using the PK area. In the flexible categorization regime this area is maximum, and the DWM can integrate a large fraction of stimulus fluctuations (Figure 2f). Indeed, we have shown that in this regime, the spiking network model, built of neural units with time-constants of 20 ms, could generate transitions by integrating fluctuations over hundreds of milliseconds (Supplementary Figure S1b). Further work would be required to quantitatively characterize the emergence of this slow integration time-scale. The PK area however, is not a measure of accuracy, when accuracy is defined as the ability to discriminate the sign of the mean stimulus evidence, μ. Thus, the accuracy in the DWM is maximal for *σ*_*S*_≈0 (Figure 3b) but the area is close to zero (Figure 2f). This mismatch simply reflects that, in the absence of internal noise, the task does not require integrating the stimulus fluctuations. However, if we only considered stimuli with *μ*=0 and we defined the stimulus category based on the sign of stimulus integral, the accuracy would be strongly correlated with the PK area and it would be maximal in the flexible categorization regime.

Finally, equipped with the theoretical results on the attractor model, we have revisited two psychophysical studies seeking for signatures of attractor dynamics. With the data from the first study ^34^, we have tested a key prediction of the attractor models and have shown that the DWM can readily fit the crossover from primacy, to flexible categorization, to recency observed as stimulus duration increases. This fit shows that the behavioral data in this task is consistent with the presence of transitions between attractor states during the perceptual categorization process (Figure 6). We used psychophysical data from a second study ^35^, to show that in a regime close to the flexible categorization the DWM could fit the categorization accuracy as a function of stimulus strength for all memory periods (Figure 7). Thus, the described asymmetry between correcting and error transitions allowed the DWM to combine evidence from the two pulses and yield a higher accuracy than a single pulse, just like subjects did (Figure 7b, compare single vs two pulse trials using the same coherence, e.g. 6.4% vs. 6.4% + 6.4%). Models that assume perfect integration of evidence can generally store a parametric value in short-term memory but they are susceptible to undergoing diffusion over time, causing a drop in memory precision as the delay increases ^55,56^. In contrast, the fact that the accuracy did not decrease with delay duration suggests that the information stored in memory was categorical instead of parametric ^57,58^, a feature naturally captured by the DWM (Figure 7d). To further investigate whether the stored information is categorical or parametric, we propose an experiment that combines electrophysiology with psychophysics to qualitatively distinguish between these two alternatives (see Supplementary Figure S3). An alternative version of the DDMA model where the sensitivity to the second pulse was larger than to the first one could also account for the combination of the two pulses ^40^. This feature captured the slight recency effect found in the data, but it left unanswered the key question of why the subjects did not use their maximum sensitivity during the first pulse. In total, our findings provide evidence that an attractor model, working in the flexible categorization regime, can capture aspects of the data that were previously viewed as incompatible with its dynamics, and propose a series of testable predictions that may further shed light onto the brain dynamics during sensory evidence integration.

## Methods

### Model simulations

For all simulations, we solve the diffusion equation 2 using the Euler method: 

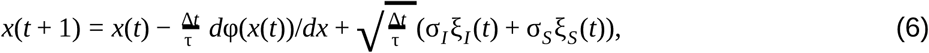

with Δ*t* = τ /40. The time constant tau of the DWM was chosen to be 200 ms to represent the effective integration time constant that emerges from the dynamics of a network ^20^. In Table 1, we summarize the parameters used in each figure.

**Table 1:** Simulation parameters for the double well and the canonical models.

| | $\mu$ | $\alpha$ | $\tau$ | $\sigma_I$ | $T$ | Bound |
| --- | --- | --- | --- | --- | --- | --- |
| Figure 1 | 0 | - | 200 ms | 0.1 | 1000 ms | 0.5 |
| Figure 2 | 0 | 1 | 200 ms | 0.1 | 1000 ms | - |
| Figure 3 DWM | 0.15 | 1 | 200 ms | 0.0 | 2000 ms | - |
| Figure 3 DDMs | 0.05 | - | 200 ms | 0 | 2000 ms | 0.5 |
| Figure 4 DWM | 0 | 1 | 200 ms | - | 1000 ms | - |
| Figure 4 DDMs | 0 | - | 200 ms | 0.08 | 1000 ms | 0.5 |
| Figure 6 | - | 0.8 | 200 ms | 0.3 | - | - |
| Figure S1 | 0 | 1 | 200 ms | 0.1 | - | - |
| Figure S2 | - | 1 | 200 ms | 0.1 | 2000 ms | - |

In Figure 4, we use stimuli with exactly zero integrated evidence, ∫*S*(*t*)*dt* = 0. For each stimulus *i*, we first created a stream of normal random variables *y*_*i*_(*t*). Then we z-score *y* and we multiplied by σ_*S*_ : 

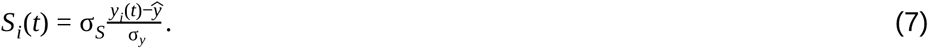

After this transformation, the mean and standard deviation of *S*_*i*_ are exactly 0 and σ_*S*_ respectively.

### Psychophysical kernel

We measure the impact of stimulus fluctuations during the course of the trial on the eventual decision by means of the so-called psychophysical kernel (PK). Put simply, given a fixed mean signal, some stimulus realizations may favor a rightward choice (say a positive decision variable) and others a leftward one. If this is the case, and we sort the stimuli over many trials by decision, we will see a clear separation which can be quantified via a ROC analysis. Mathematically, for each trial *i*, we subtract the mean evidence (μ_*i*_) of each trial *s*_*i*_(*t*) = μ_*i*_ + σ_*s*_ξ_*I*_ to avoid that the distributions of stimuli that produce left and right choices are trivially separated by their mean evidence: 

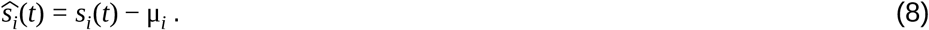

Thus 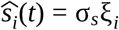 are simply the stimulus fluctuations. Then, for each time *t*, we compute the probability distribution function of the stimuli that produce a right 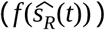 or left 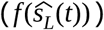 choice. The PK is the temporal evolution of the area under the ROC curve between these two distributions 

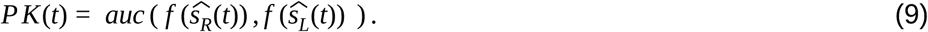

### Normalized psychophysical kernel area and primacy-recency index

In order to quantify the magnitude and the shape of a PK, we defined two measures, the PK area and the PK slope:

1. The normalized PK area is a measure of the overall impact of stimulus fluctuations on the upcoming decision, it ranges from 0 (no impact) to 1 (the stimulus fluctuations are perfectly integrated to make a choice). It is defined as 

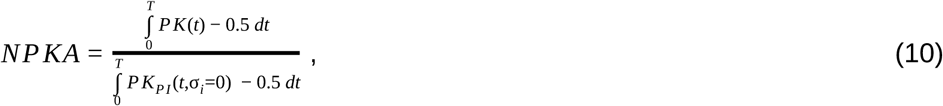

where *T* is the stimulus duration. *NPKA* is the PK area normalized by the PK area of a perfect integrator in the absence of internal noise (σ_*i*_ = 0), i.e. an ideal observer.
2. The normalized PK slope is the slope of a linear regression of the PK, normalized between −1 (decaying PK, primacy) to +1 (increasing PK, recency). Because we wanted the PK slope to quantify the shape of the PK rather than its magnitude (which is captured by the PK area), we first normalized the PK to have unit area, 

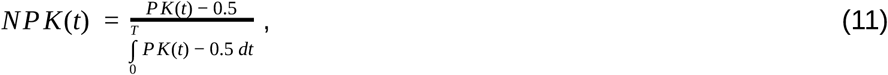

where *T* is the stimulus duration. We then fit the NPK with a linear function of time, 

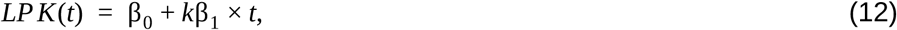

where β_1_ is the PK slope and 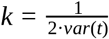 is a factor that normalizes the PK slope to the interval (−1, +1).

### Accuracy for the double well model

To compute the accuracy for the double well model (DWM), we assume that the time spent in the unstable region is much shorter than the time spent in one of the attractors. This assumption allows us to treat the system as a Continuous Markov Chain (CMC) with only two possible states correct and error. The first step is to compute the probability of first visiting the correct attractor which will be used as the initial state of the CMC (Gardiner 1985) 

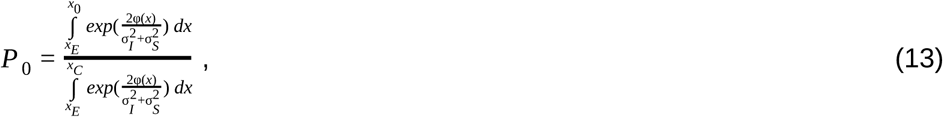

where φ is the potential in equation 3, *x*_*C*_ and *x*_*E*_ are the *x* values of the correct and error attractors whereas *x*_0_ = 0 is the initial position of *x*. The integrals of *P* _0_ can be computed assuming that the term *x*4 is very small for values of *x*_0_ ≃ 0 : 

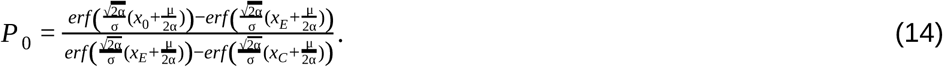

The second step is to compute the correcting and error transition rates ^37,59^ 

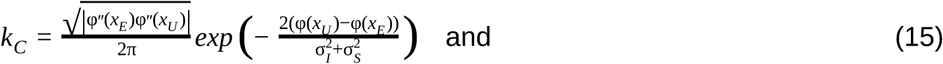

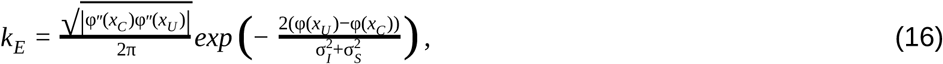

where *x*_*U*_ is the *x* position at the unstable state. These are the transition rates of a Continuous Markov Chain with only two states: correct and incorrect. The probability of making a correcting and error generating transition during a trial are ^60^: 

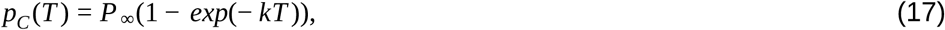

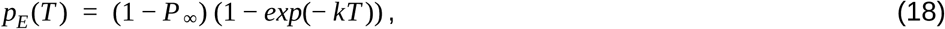

Where *k* = *k*_*C*_ + *k*_*E*_, *T* is the stimulus duration and 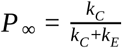 is the probability of the stationary state being the correct one (*T* → ∞). Finally, the probability of being in the correct attractor given the model and stimulus parameters is 

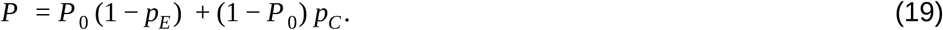

The probability of correct is the probability to first visit the correct attractor and remain in it (*P* _0_ (1 − *p*_*E*_)) plus the probability to first visit the error attractor and correct the initial decision ((1 − *P* _0_)*p*_*C*_). To be more quantitative, we can compute the ratio between the probability of a correcting (equation 17) and a error-generating transition (equation 18): 

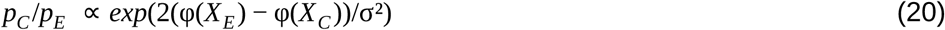

For small values of the mean signal μ << 1, we can rewrite the ratio between the correcting and error-generating transitions as a function of the potential parameters. To this aim we compute the fixed points of order ϑ(ε^2^) using 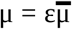 where 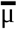 is a parameter of order 1 and *x* = *x*_0_ + ε*x*_1_ : 

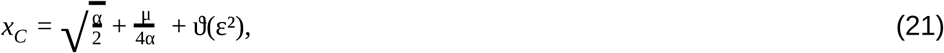

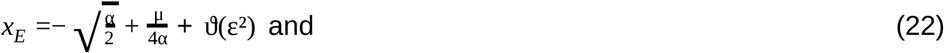

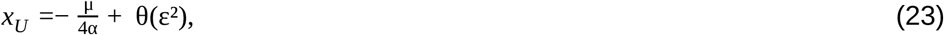

where *x*_*U*_ is the *x* position of the unstable state (note that *x*_*U*_ = 0 when μ = 0) and *x*_*C*_ (*x*_*E*_) is the position of the correct (error) attractor. Using these fixed points, the ratio between the correcting and error-generating transitions is 

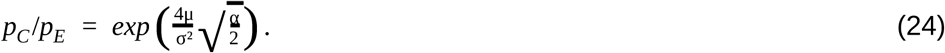

Which shows that the ratio between correcting transitions and error-generating ones increases exponentially with the mean stimulus (μ) as long as stimulus fluctuations are not too large. These probabilities are illustrated in Figure 3c, *p*_*C*_ increases steeply as a function of stimulus fluctuations even before *p*_*E*_ reaches non-negligible values and for large stimulus fluctuations both probabilities tend to 0.5.

### Compatible parameters with a non-monotonic accuracy

Here we investigate the parameter range in which the accuracy is non-monotonic with the stimulus fluctuations. Concretely, we compute the critical values of the mean stimulus evidence (μ) and the internal noise (σ_*I*_) beyond which the performance decays monotonically with the stimulus fluctuations σ_*S*_. Plugging the attractor positions for weak stimulus strength μ from equations 21, 22 and 23 into equations 15 and 16, the transition rates are 

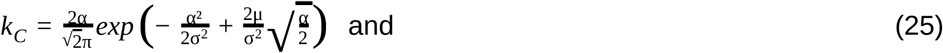

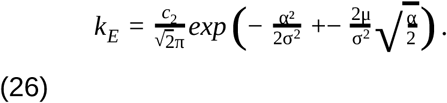

We define the total level of noise (σ^2^) as the sum of the stimulus fluctuations and the internal noise, 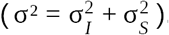. If there is a non-monotonicity of the accuracy with σ, we should find a maximum of the probability of correct when the error attractor was first visited (equation 17). 

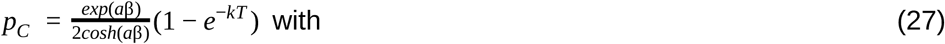

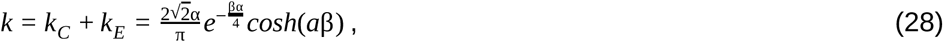

where 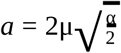 and 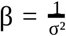. To check the existence of a local maximum we take the derivative of *p*_*C*_ with respect to σ 

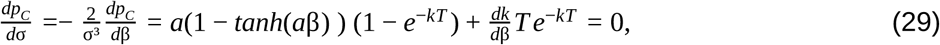

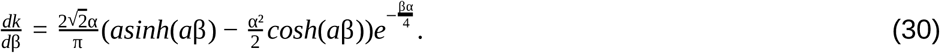

For small values of μ, the arguments of the trigonometric hyperbolic functions are very small and they can be approximated by *sinh*(*a*β) ≃ 0, *tanh*(*a*β) ≃ 0 and *cosh*(*a*β) ≃ 1. Using these approximations, equations 29 and 30 can be simplified as 

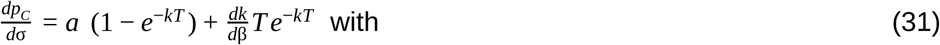

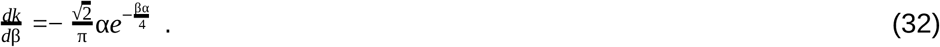

For small values of σ, 1 − *e*^−*kT*^ ≃ 0. However there is always a large enough *T* so that 1 − *e*^−*kT*^ ≃ 1. The local maximum of the accuracy must be in the region where these two effects are of the same order *kT* ∼ θ(1) and the two terms in equation 31 cancel each other. Thus we defined 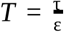 and 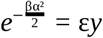, plugging these into equation 31, we obtain

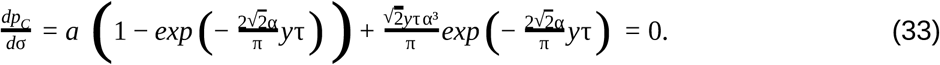

Let us define 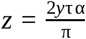. To have a maximum of *p*_*C*_, there must be a solution to the following implicit equation 

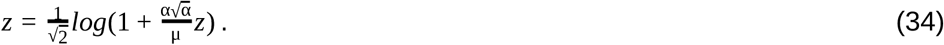

Using the definitions of τ, *y* and *z*, we find a maximum of the probability of *p*_*C*_ as a function of the solution (*z*_0_) of the implicit equation 34: 

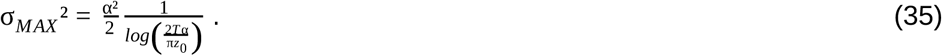

To find the maximum of the accuracy, we derive equation 19 respect to σ : 

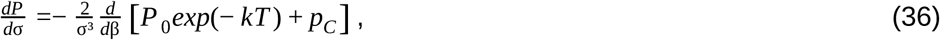

where we rewrite equation 19 as a function of *p*_*C*_ and *P* _0_. As long as 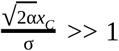, the probability to first visit the correct attractor (equation 14) is well approximated by 

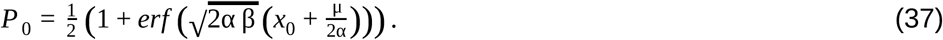

In the range of parameters where μ is small and β is large we can assume that βμ^2^ << 1, 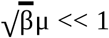 and μ β >> 1. Using these inequalities and the definition of *P* _0_ given in equation 37, we can simplify equation 36 to 

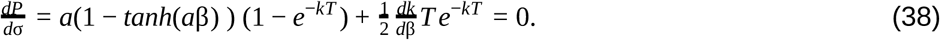

With these simplifications, the derivative 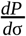 is equivalent to the derivative 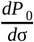 (equation 31) with a 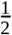 factor in the second term. This factor modifies the implicit equation 34 to 

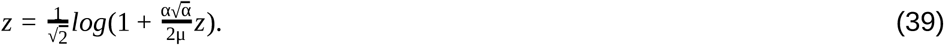

Then using the definition of *z*, the critical value of the internal noise for which accuracy decreases monotonically with the stimulus fluctuations is 

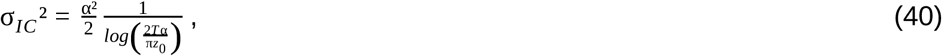

where *z*_0_ is a solution of the implicit equation 39. This implicit equation has two solutions, the trivial solution *z*_0_ = 0 when σ is small and there are no transitions and a positive solution *z*_0_ > 0. The accuracy is non-monotonic with the stimulus fluctuations when the positive solution exists. The positive solution of a general implicit equation of the form (*x* = *log*(1 + *cx*)) exists when the derivative of the right term at *x* = 0 is larger than 1, (*c* > 1). In the case of equation 39, the positive solution exists when 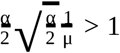. Thus there is a critical value of the mean evidence (μ) above which the accuracy decreases monotonically with the stimulus fluctuations 

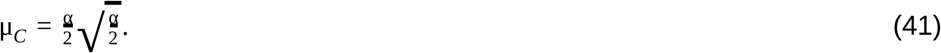

### Spiking network

We consider a network of recurrently coupled integrate-and-fire neurons, similar to ^28^. The network consists of two populations of excitatory neurons (A and B), both of which are recurrently coupled between them and to a population of inhibitory interneurons (I). We study the case in which the system is near a steady bifurcation to a winner-take-all state. It is in the vicinity of the bifurcation that the dynamics of the network can be captured in a one-dimensional amplitude equation which describes the slow evolution along the critical manifold ^28^. The evolution of the membrane potential 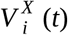 from the *i*-th neuron in population *X* is given by: 

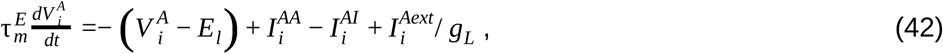

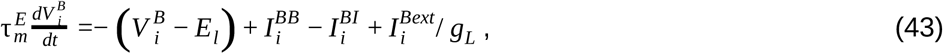

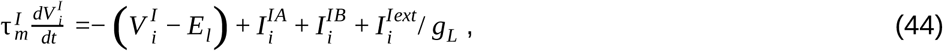

where the synaptic input voltages of the form *I*_*XY*_ indicate interactions from neurons in population *Y* to neurons in population *X*, while external synaptic inputs are given by *I*^*Xext*^.

The synaptic inputs are sums over all postsynaptic potentials (PSPs), modeled as exponential functions with a delay. The synaptic inputs take the form 

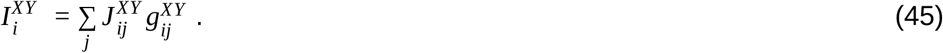

The dynamics of excitatory and inhibitory synapses are described by 

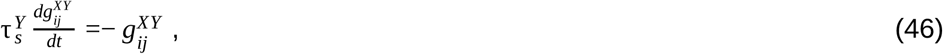

After the presynaptic neuron *j* fires a spike at time 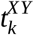 the corresponding dynamic variable is incremented by one at 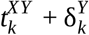, that is after a delay 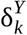.

External synapses have instantaneous dynamics 

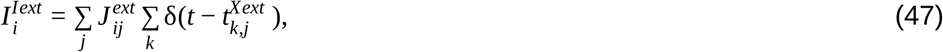

i.e. a presynaptic action potential from neuron *j* of the external population at time 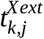 results in an instantaneous jump of the external synaptic input variable. A spike is emitted whenever the voltage of a cell from an excitatory (inhibitory) population crosses a value Θ, after which it is reset to a reset potential *V* _*r*_.

We consider the case of sparse random connectivity for which, on average, each neuron from population X receives a total of *C*_*XY*_ synapses from population Y. The pairwise probability of connection is thus ε _*XY*_ = *C*_*XY*_ /*N*_*Y*_, where *N*_*A*_ = *N*_*B*_ = *N*_*E*_ and *N*_*I*_ are the number of neurons in the respective populations. For nonzero synapses we choose 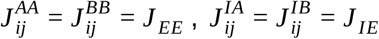 and 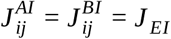.

The stimulus input current is modeled similar to ^24^, with the exact same stimulus input being injected to each neuron in each of the two excitatory populations. The stimulus input onto each of the excitatory populations A and B is given by 

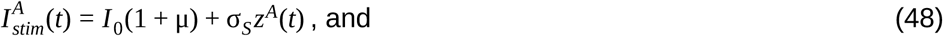

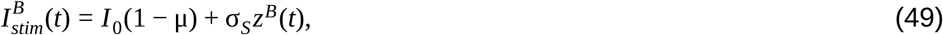

where the first term describes the mean stimulus input onto each population and the second term the temporal modulations of the stimulus with standard deviation σ_*stim*_. The term μ parametrizes the mean difference of the two stimulus inputs and it captures the amount of net stimulus evidence favoring one choice over the other (i.e. μ =0 represents an ambiguous stimulus with zero mean sensory evidence). Finally, *z*^*A*^(*t*) und *z*^*B*^ (*t*) are independent realizations of an Ornstein-Uhlenbeck process, defined by 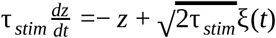, where ξ(*t*) is Gaussian white noise (mean 0, variance dt).

**Table 2:**
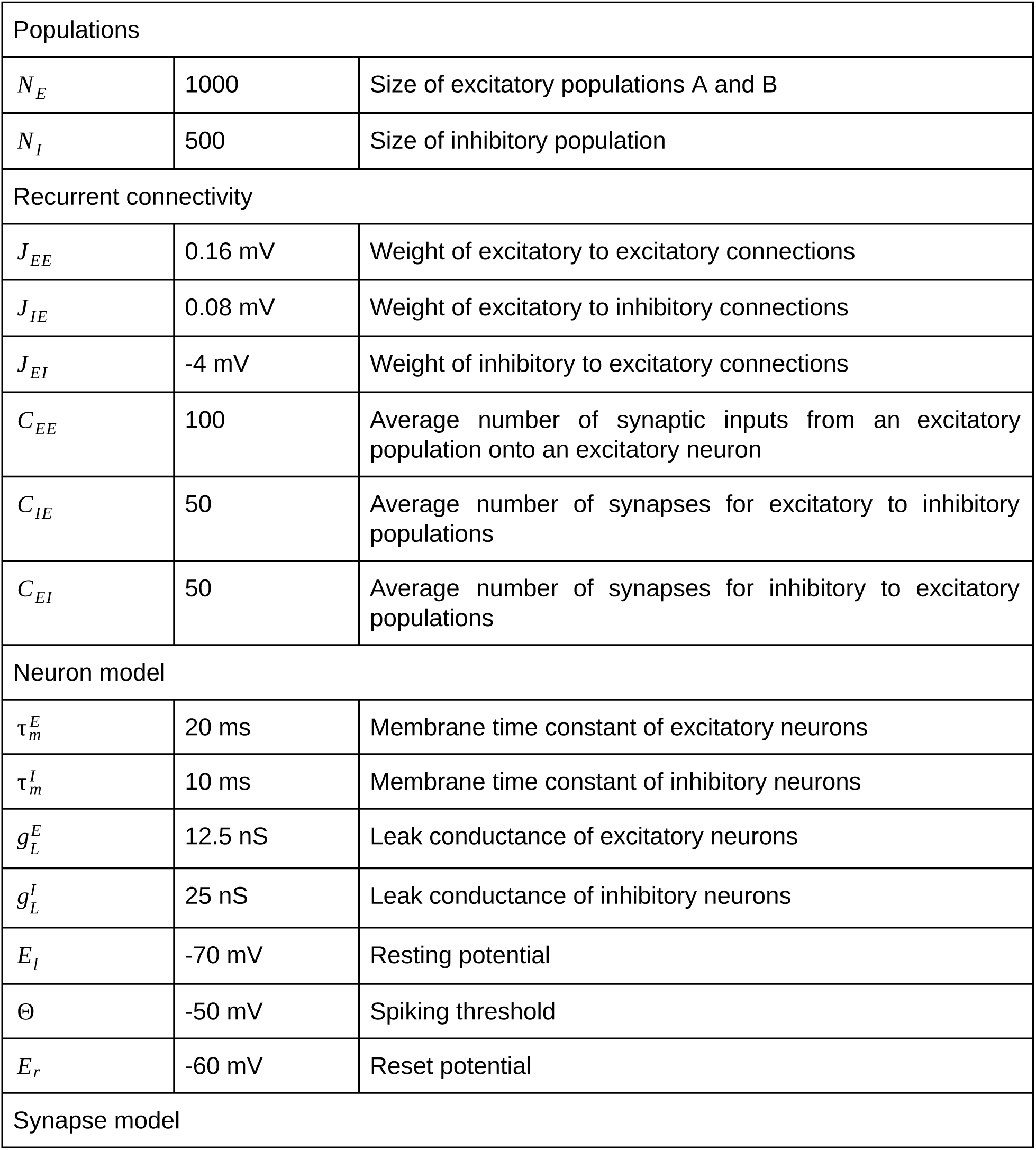
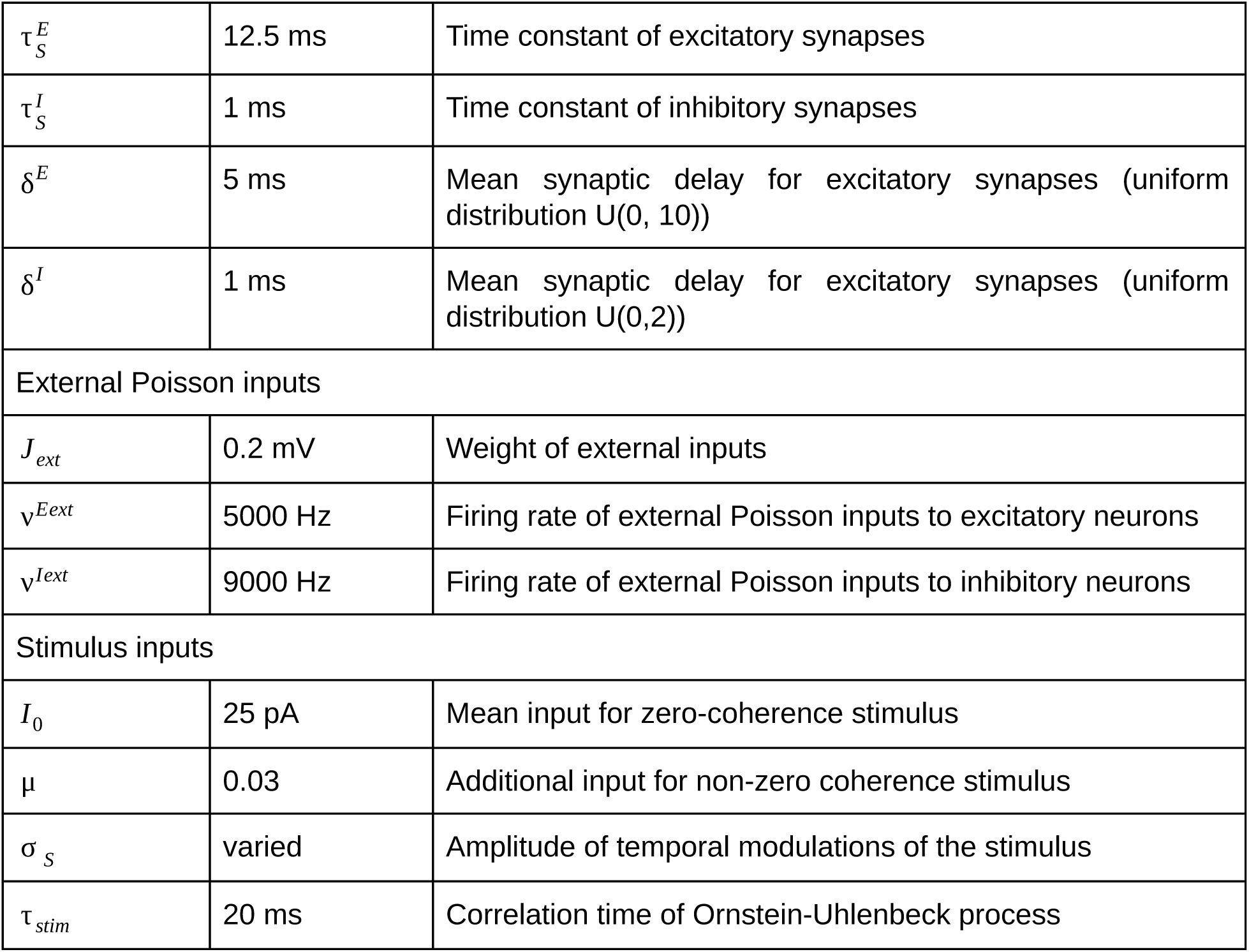
Simulation parameters for the spiking neural network model.

### Psychophysical data and model fitting

In Figure 6, we used data from experiments 1 and 4 from ^34^ with a total of *N* = 21 humans subjects (*N* = 13 in experiment 1 and *N* = 8 in experiment 4). The data can be accessed here: https://doi.org/10.1371/journal.pcbi.1004667. The stimuli consisted of two brightness-fluctuating round disks. In each stimulus frame (duration 100 ms), the brightness level of each disk was updated from one of two generative Gaussian distributions that had the same variance but different mean: either one distribution had a high mean value and the second a low value or vice versa. At the end of the stimulus, the subjects had to report the disk with a higher overall brightness (i.e. which disc corresponded to the generative distribution with higher mean). Incorrect responses were followed by an auditory feedback. Trials were separated into 5 equal length segments, in 80 % of the trials, a congruent or incongruent pulse of evidence was presented at a random segment. This increase or decrease of evidence was corrected in the rest of the segments and as a consequence the stimuli were anticorrelated. In experiment 1 stimuli with 1,2 or 3 seconds duration were presented in blocks of 60 trials whereas in experiment 4, the stimulus duration was 5 seconds. We computed the PK using the procedure described above (see section Psychophysical kernel) but first computing the difference in brightness of the two disks. We also subtracted the mean difference in order to have a one-dimensional stimulus trace with zero mean. Namely 

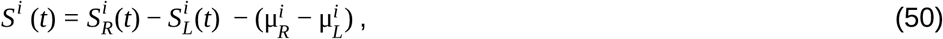

where 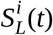 is the brightness of the *t*-th frame of the left disk during the *i*-th trial and 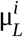 is the mean of the generative Gaussian distribution for the left disc in the i-th trial. We computed the PKs standard error of the mean using bootstrap with 1000 repetitions.

To compute the PK of the DWM we simulated equation 6 using stimuli with the exact same temporal fluctuations in evidence than the stimuli presented to the subjects. We modeled it by updating μ^*i*^ (*t*) from equation 3 with the difference in brightness at each time between the right and left disk:

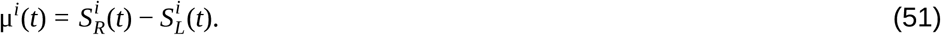

Note that in this framework the stimulus fluctuations were set to zero σ_*S*_ = 0 because σ_*S*_ was captured inside μ^*i*^(*t*). The DWM parameters (α =− 0.8, σ_*I*_ = 0.3 and τ = 200 *ms*) were tuned to account for the change from primacy to recency with the stimulus duration.

### Primacy-recency index for the two pulses trials

In Figure 7, we define the primacy-recency index 

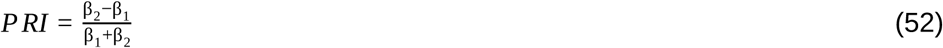

where β_1_ and β_2_ are the coefficients of a logistic regression with the coherence of the first and second pulse as predictors: 

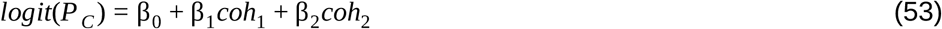

Similar to the Normalized PK slope, the primacy-recency index ranges from −1 (primacy) to 1 (recency).

### Double well model fitting

In Figure 7, we use data from two studies performing the same experiments (Kiani et al 2013 and Tohidi-Moghaddam 2018). We extract the accuracy of the subjects directly from the paper figures (with GraphClick, a software to extract data from graphs) and the number of trials from the methods of the papers. We pool the data from the two experiment and we compute the mean accuracy in each condition *i* as 

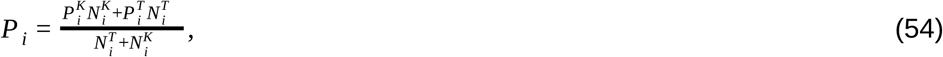

where N_i_ is the number of trials in condition *i*, the data with superindex *K* and *T* were extracted from ^35^ and ^40^ respectively. The 95% confidence interval of *P* _*i*_ is: 

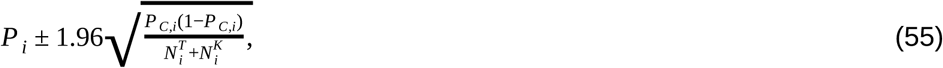

In these experiments, the human subjects had to discriminate between left and right motion direction of a random dots stimulus. The experimenters interleaved trials with one and two pulses of 120 ms. For single pulse trials the possible coherence levels were 0%, 3.2%, 6.4%, 12.8%, 25.6% and 51.2%. For double pulse trials, the pulses were separated by a delay of 0, 120, 360 or 1080 ms and the coherences were randomly chosen from 3.2%, 6.4% and 12.8% (nine different coherence sequences). In both papers, they reported that the subjects’ accuracy in double pulses trials was independent of the delay. Thus we assume that, in the DWM, the internal noise was too small to drive transitions during the delay and we pool the data across delays to compute the accuracy for each coherence sequence. We fit the model by maximizing the log-likelihood (Nelder-Mead algorithm): 

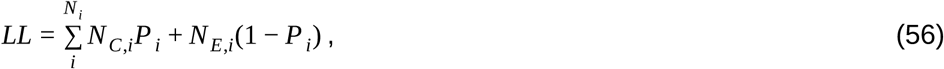

Where *NC*,*i* and *N*_*E*,*I*_ are the number of correct and error trials for each coherence sequence *i* whereas *P* _*I*_ is the accuracy for sequence *i* predicted by the DWM.

For single pulse trials, we computed *P* _*i*_ as 

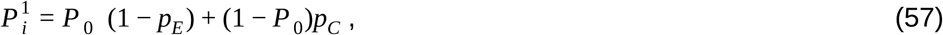

where *P* _0_, *p*_*C*_, and *p*_*E*_ were computed using equations 14, 17 and 18 whereas the super index indicates the pulse number. Note that we are assuming that the time spent for the decision variable in the unstable state is short compared with the pulse duration. In this model, the decision variable starts in the correct attractor with probability *P* _0_. Similarly for double pulse trials the probability of correct is: 

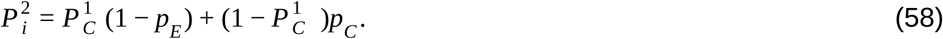

The potential and the diffusion equation can be written as 

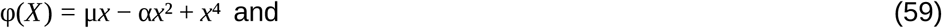

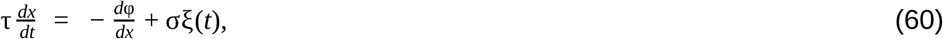

where μ is a linear scaling of the coherence to *x* units (μ = *kcoh*) and σ represents the two sources of noise, the internal noise and the stimulus fluctuations 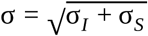. The two sources of noise can not be fitted separately because the only difference between them is that the internal noise is also activated during the delay (Figure 7a). But internal noise does not have any impact during the delay. Thus it is impossible to distinguish σ_*I*_ in the range 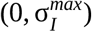 where 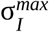 is the maximum σ_*I*_ without transitions during the delay. For this reason, we assume that there are no transitions during the delay and we only fit the total noise σ. The parameters that maximize equation 56 and their 95% confidence interval are *k*^*^ = 0.012 ± 0.0011, α* = 0.70 ± 0.05, σ* = 0.52 ± 0.05 and τ * = 3.3 ± 0.5. To compute the confidence intervals, we assume that the likelihood function around the best-fit parameters is a multi-dimensional Gaussian. Then the confidence intervals are two times the diagonal of the inverse of the Hessian matrix ^18,61^. The Hessian matrix is the matrix of second derivatives and we compute it numerically using the finite difference method.

Although we can not fit the internal and the stimulus sources of noise separately, we can study the range of internal noise 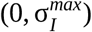 that produces a negligible number of transitions (< 1%) during the delay (up to 1 s) and thus is compatible with the psychophysical data. For the parameters that maximize the likelihood this range is (0, 0.32), indicating that the DWM is robust to perturbations during the delay even when the magnitude of the internal noise represents a substantial part of the total noise 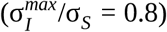 (Figure 7d). We also compute the accuracy of the perfect integrator as a function of the delay (Figure 7d). To be able to compare both models, we adjust the scaling factor of the evidence to match subjects’ accuracy for the shortest delay (μ_*PI*_ = 0.44*kcoh* where *k* is the scaling of the DWM), and we use the parameters τ, σ_*S*_ and σ_*I*_ that maximize the DWM.

## Data availability

Data shown in figure 6 can be accessed here: https://doi.org/10.1371/journal.pcbi.1004667. The data shown in figure 7 was extracted directly from the manuscript using GraphClick ^35,40^.

## Code availability

The code to generate the figures of the paper will be uploaded in github.

## Acknowledgements

We thank Tobias H. Donner and Niklas Wilming for excellent discussions. The research leading to these results has received funding from “la Caixa” Foundation (to G.P.O.), the Spanish Ministry of Economy and Competitiveness together with the European Regional Development Fund (RYC-2015-17236 and BFU2017-86026-R to K.W, MTM2015-71509-C2-1-R and RTI2018-097570-B-I00 to A.R. and SAF2015-70324-R to J.R.) and from the Generalitat de Catalunya (grant AGAUR 2017 SGR 1565 to A.R., J.R. and K.W.). This work has received funding from the European Research Council (ERC) under the European Union’s Horizon 2020 research and innovation programme (ERC-2015-CoG - 683209 PRIORS to J.R). Part of this work was developed at the building Centre Esther Koplowitz, Barcelona.

## Author contributions

All the authors contributed to the design of the study and to the interpretation of the results. G.P.O. performed the simulations and the analysis of the canonical and the double well models. G.P.O. and A.R. derived analytical expressions from the DWM. K.W. and G.P.O. performed the simulations and analysis of the spiking network. All the authors wrote the paper.

## Supplementary Information

**FIgure S1.**
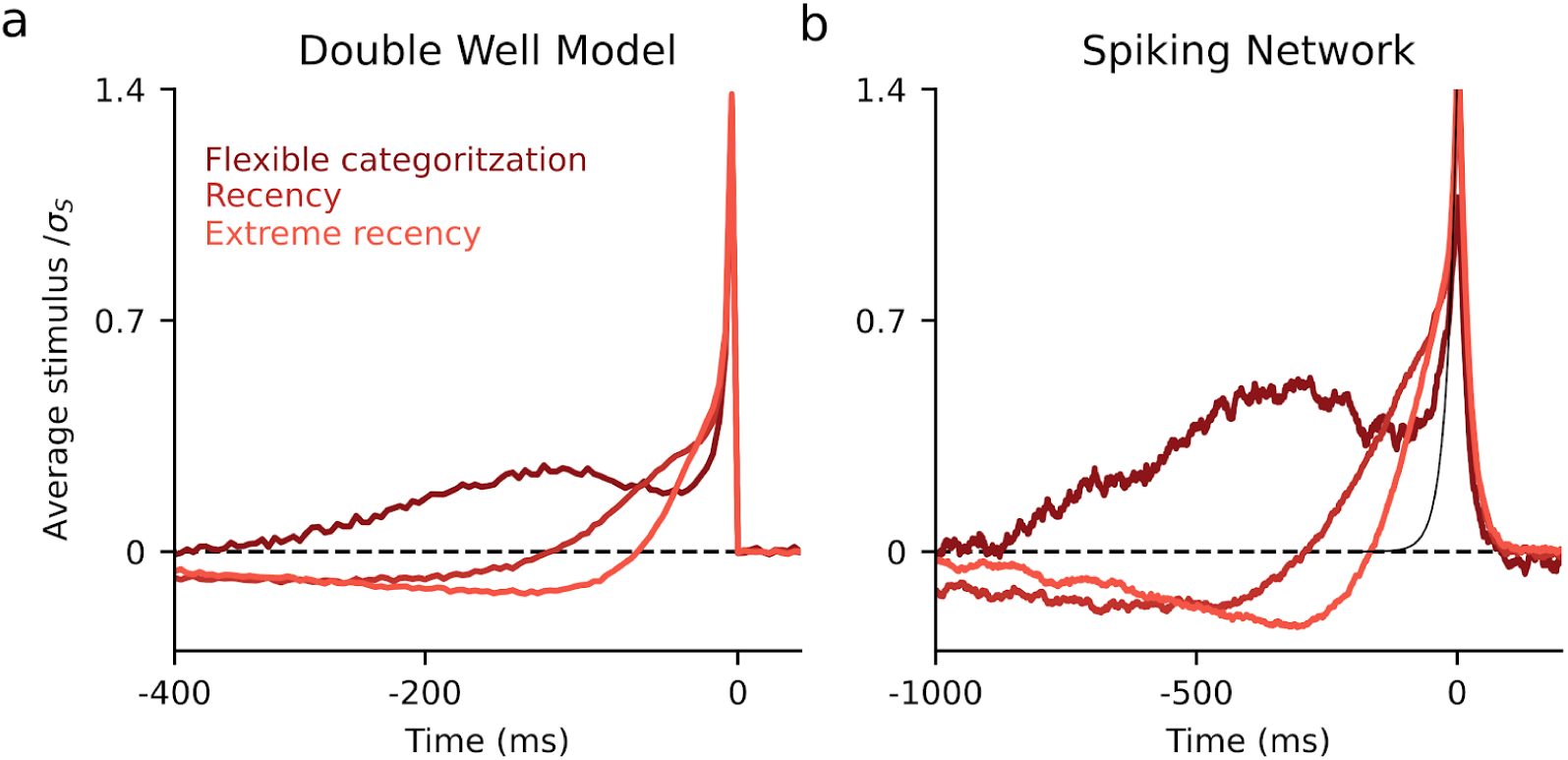
Stimulus integration during a transition between attractor states. **(a)** Average stimulus aligned to the transition time (t = 0 ms) in the flexible categorization and recency regime for the DWM (σ_S_ = 0.5, 1 and 1.5) We introduced two thresholds at the attractor states 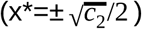 and we defined a transition when the decision variable reached one of the thresholds when the opposite had been previously reached. The peak at t=0 is an artifact produced by crossing the threshold (i.e the last fluctuations was always in favour to cross the threshold). In the flexible categorization regime, the transitions occur when the stimulus favours them for hundreds of ms, considerably longer than the time constant of the system (τ = 200 *ms*). Thus the integration of the stimulus continues even when an attractor state has been reached. As stimulus fluctuations increase, the transitions become faster and the system moves into a regime where the transitions are based on momentary evidence rather than an integration of the stimulus (extreme recency). **(b)** Same as in (a) for the spiking network with σ_S_=4, 9 and 13, and thresholds at r_A_-r_B_=±25Hz where r_A_ and r_B_ are the firing rates of the two excitatory populations. The stimulus was taken as the difference of the input to populations A and B, 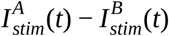 and *μ*=0. While the dynamics of the DWM is governed by a single time constant *τ*, in the spiking network several time constants contribute to the dynamics (Table 2; membrane time constants 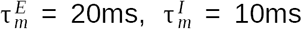, synaptic time constants 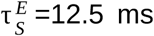, 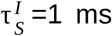, synaptic delays). Moreover, the stimulus fluctuations are not white noise but correlated noise, realized as an Ornstein-Uhlenbeck process with *τ*_stim_=20 ms. The expected decay of the average stimulus is given by the stimulus autocorrelation (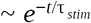 ; black line). Despite all these factors, the transition-triggered average stimuli are qualitatively similar to the DWM. In particular, the longest integration occurs in the flexible categorization regime and exceeds by far all the intrinsic time constants of the network.

**Figure S2.**
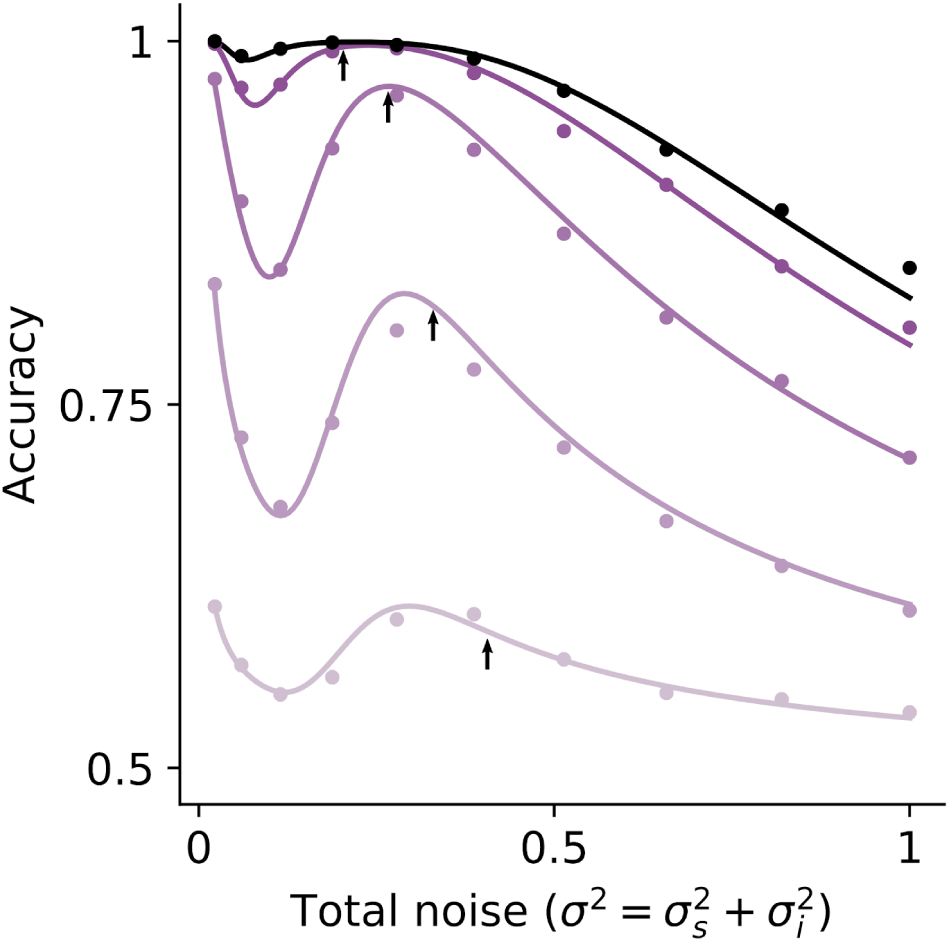
Critical internal noise and mean stimulus evidence compatible with the non-monotonic relation between accuracy and stimulus fluctuations. Accuracy versus the total noise, 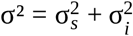 obtained from simulations (dots) and theory (equation 19, solid line) for different mean stimulus evidence μ = 0.03, 0.1, 0.2 and 0.3 from light to dark purple as well as the for the critical value μ_*C*_ = 0.35 in black. The bump in accuracy occurs if the probability of a correcting transition given by the second term in equation 19 is large enough when the error transition are not activated (1 − *p*_*E*_ ≃ 1). In other words, the accuracy decrease monotonically with σ^2^ if in the limited regime where there is a large asymmetry between correcting (*p*_*C*_) and error (*p*_*E*_) transitions (Figure 3b), the probability of an error initial categorization (1-*P*_*0*_) is small and the number of correcting transitions is negligible. Note that the bump becomes smaller when μ → 0. Thus, intermediate values of mean stimulus evidence are recommended to experimentally test this non-monotonic relation. The black arrows indicate the critical value of the internal noise for a non-monotonic relation between the accuracy and the stimulus fluctuations from equation 40. It is precisely the value of the local maximum in the total noise.

**Figure S3.**
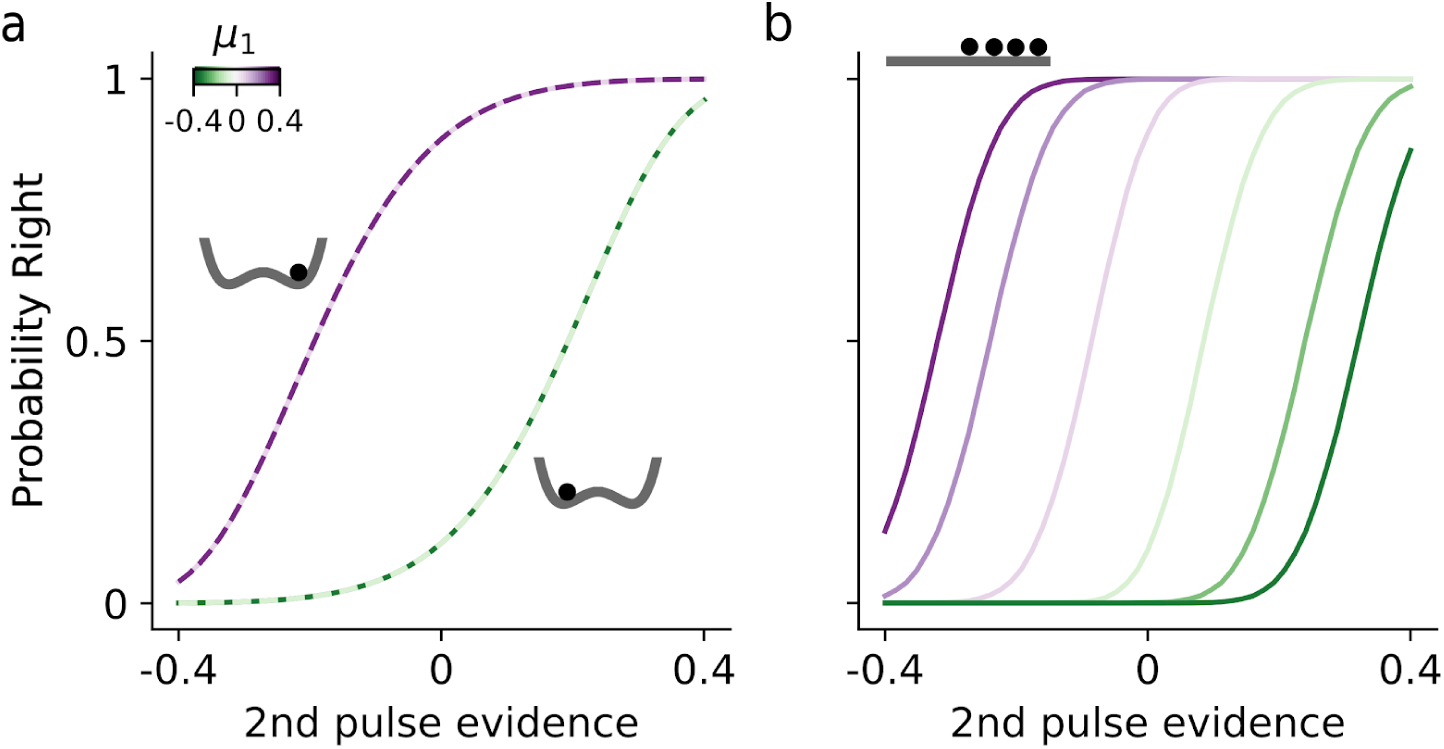
Categorical versus parametric working memory. To further illustrate the implications of the DWM categorization dynamics in comparison to a perfect integrator, in which intermediate values of the decision variable are meta-stable, we propose a modification of the two pulses experiment in which using invasive or non-invasive neurophysiological recordings, the decision variable during the delay period could be read-out. The aim of this experiment would be to investigate if the information about the first pulse stored during the delay is a categorical (DWM) or a parametric (Perfect Integrator) value. We would use the sign of the decision variable read out during the delay to sort the trials between correct and incorrect initial categorization. Then, using only the correct initial categorized trials, we could plot the accuracy of the final response as a function of the second pulse evidence. (**a**) During the delay, the DWM categorically stores the initial categorization, thus, the probability to choose right is independent of the strength of μ_1_. Note that because we only consider correct initial categorization trials, purple (green) lines represent those trials where the decision variable was in the right (left) attractor during the delay (see inset). (**b**) In contrast, for the perfect integrator, the information stored during the delay is a parametric value proportional to μ_1_ (see inset) and thus the decision depends on the strength of μ_1_. Given that this experiment requires to read-out the decision variable during the delay, one could think that it should be enough to directly assess whether the distribution of the decision variable during the delay is categorical or parametric. However, it has been shown that different brain areas encode the decision variable with different degrees of categorization ^62^. Thus the result could depend on the brain area used to decode the decision variable. In contrast, in our experimental paradigm, we would assess whether subjects are indeed using a categorical or parametric representation of the first pulse independently of what can be decoded in different brain areas.

